# MyD88-Dependent Signaling Drives Toll-Like Receptor-Induced Trained Immunity in Macrophages

**DOI:** 10.1101/2022.08.23.504963

**Authors:** Allison M. Owen, Liming Luan, Katherine R. Burelbach, Margaret A. McBride, Cody L. Stothers, Olivia A. Boykin, Kalkena Sivanesam, Jessica F. Schaedel, Tazeen K Patil, Jingbin Wang, Antonio Hernandez, Naeem K. Patil, Edward R. Sherwood, Julia K. Bohannon

## Abstract

Immunocompromised populations are highly vulnerable to developing life-threatening infections. Strategies to protect patients with weak immune responses are urgently needed. Employing trained immunity, whereby innate leukocytes undergo reprogramming upon exposure to a microbial product and respond more robustly to subsequent infection, is a promising approach. Previously, we demonstrated that the TLR4 agonist monophosphoryl lipid A (MPLA) induces trained immunity and confers broad resistance to infection. TLR4 signals through both MyD88- and TRIF-dependent cascades, but the relative contribution of each pathway to induction of trained immunity is unknown. Here, we show that MPLA-induced resistance to *Staphylococcus aureus* infection is lost in MyD88-KO, but not TRIF-KO, mice. The MyD88-activating agonist CpG (TLR9 agonist), but not TRIF-activating Poly I:C (TLR3 agonist), protects against infection in a macrophage-dependent manner. MPLA- and CpG-induced augmentation of macrophage metabolism and antimicrobial functions is blunted in MyD88-, but not TRIF-KO, macrophages. Augmentation of antimicrobial functions occurs in parallel to metabolic reprogramming and is dependent, in part, on mTOR activation. Splenic macrophages from CpG-treated mice confirmed that TLR/MyD88-induced reprogramming occurs in vivo. TLR/MyD88-triggered metabolic and functional reprogramming was reproduced in human monocyte-derived macrophages. These data show that MyD88-dependent signaling is critical in TLR-mediated trained immunity.

## INTRODUCTION

Despite medical advances, infections frequently progress to the life-threatening clinical condition of sepsis, especially in immunocompromised patients. As evidenced by the ongoing COVID-19 pandemic, patients with advanced age or underlying co-morbidities such as chronic lung diseases, cancer, cardiovascular disease, or traumatic injury are at serious risk for life-threatening infections (^1, 2^). Hospital stays increase exposure to opportunistic pathogens which immunocompromised patients are unable to fight due to insufficient immune responses. Nosocomial infections affect 2 million patients and cause more than 90,000 deaths every year in the United States (^3^). Further complicating the matter, antibiotic resistance in microbes such as *Pseudomonas aeruginosa* and *Staphylococcus aureus* remains an ever-growing threat, with the Centers for Disease Control and Prevention reporting that nearly 3 million antibiotic resistant infections occur in the US each year (^4^). The development of new antibiotics is slow, with the most recent discovered in 1987, and the investment into new antibiotic research only temporarily addresses the situation given the unfortunate reality that pathogens will evolve to evade new antibiotics (^4^). Therefore, the development of novel protection strategies is an urgent priority, and immunomodulatory therapies hold strong potential to fill this need (^5^).

Immune memory is classically thought to be a characteristic of the adaptive immune system, however recent reports show that the innate immune system also has the ability to develop memory of prior pathogen exposure. Evidence shows that innate leukocytes can be “trained” to mount a more robust immune response to subsequent infection, a phenomenon termed *trained immunity* (or *innate immune training*) (^6^). Much of the recent literature has focused on the ability of β-glucan, a component of the fungal cell wall, to induce trained immunity (^7, 8^). However, the capability of the Toll-like Receptor (TLR)-4 agonist lipopolysaccharide (LPS) to protect against subsequent infection was described as early as the 1950s (^9^). More recently, our research group has demonstrated that other TLR4 ligands, including the lipid A derivative compounds monophosphoryl lipid A (MPLA) and phosphorylated hexaacyl disaccharides (PHADs), increase leukocyte recruitment to sites of infection, protect against organ injury, and improve survival in a broad array of clinically relevant infections, and these effects extend up to two weeks (^10–15^). Our group has previously shown that macrophages are essential for the survival benefit conferred by MPLA; further, this protection is mediated, at least in part, by metabolic rewiring characterized by sustained augmentation of glycolysis and mitochondrial oxygen consumption capacities (^13, 16^).

In a ligand-specific manner, activated TLRs recruit the adapter molecules myeloid differentiation primary response differentiation gene 88 (MyD88) or Toll/IL-1R (TIR) domain-containing adapter producing interferon-β (TRIF). Recruitment of MyD88 or TRIF elicits pathway-specific downstream signaling and gene transcription events, with MyD88 signaling culminating in production of pro-inflammatory mediators, and TRIF signaling resulting in production of Type I interferons. TLR4 uniquely signals through both the MyD88- and TRIF-dependent signaling cascades, but the respective contributions of these pathways in triggering trained immunity and host resistance to infection is unknown. Other investigators have shown that the efficacy of MPLA as a vaccine adjuvant is mediated through TRIF-dependent signaling(^17^). On the other hand, our lab has found that treatment with the MyD88-selective TLR9 agonist CpG, but not TRIF-selective TLR3 agonist Poly I:C, preserves core body temperature, increases leukocyte recruitment, and improves bacterial clearance to an acute intraperitoneal infectious challenge with *Pseudomonas aeruginosa* (^12^). In vitro analysis of antimicrobial functions of bone marrow-derived macrophages (BMDMs) showed that MPLA- and CpG-treated cells exhibited increased phagocytic and respiratory burst capacity. These data suggest that MyD88, rather than TRIF, is important for driving TLR-mediated training of innate immunity.

In this study, we aimed to determine the relative contributions of MyD88- and TRIF-dependent signaling pathways and the role of macrophages in driving TLR-mediated infection resistance. Further, we sought to evaluate the role of these signaling pathways in stimulating metabolic reprogramming, a defining characteristic of trained immunity. We establish that MyD88-dependent, but not TRIF-dependent, signaling in macrophages is an essential driver of TLR-mediated trained immunity and resistance to infection. Using a combination of complimentary in vivo, in vitro, and ex vivo models, we show that macrophage metabolic rewiring is likewise MyD88-dependent and is associated with augmented antimicrobial immunity. We corroborated these findings using human monocyte-derived macrophages. These data advance current knowledge and provide insights that can be used to develop more specific TLR agonist-based immunotherapies aimed at inducing resistance to infection.

## RESULTS

### MyD88, but not TRIF, activation is required for TLR-mediated protection against systemic *S. aureus* infection

We previously showed that the TLR4 agonist monophosphoryl lipid A (MPLA) protects against a broad array of clinically relevant pathogens (^10, 11, 13^). As TLR4 triggers both MyD88- and TRIF-dependent signaling cascades, we sought to determine the relative contribution of these pathways in triggering trained immunity. Further, it was unknown whether activation of other TLRs can elicit trained immunity. To investigate these questions, wild type, MyD88-KO and TRIF-KO mice were treated with TLR agonists (20 µg *i.v.*) or vehicle for two consecutive days and were inoculated with *S. aureus* the following day (**Figure 1A**). Consistent with our previous findings, MPLA-treated wild type mice had significant survival benefit (80% survival) compared to vehicle-treated controls (**Figure 1B**). MPLA did not confer survival benefit in MyD88-KO (0% survival, **Figure 1C**) but was effective in TRIF-KO mice (100% survival, **Figure 1D**). The minimum lethal dose of *S. aureus* was adjusted for MyD88-KO mice due to increased sensitivity to infection, which resulted in similar blood bacteria loads in vehicle-treated mice from all three strains (**Figure 1E**). This was associated with a trend of decreased blood bacterial burden in MPLA-treated wild type and significantly decreased bacterial burden in TRIF-KO, but no difference in bacterial burden in MyD88-KO, mice.

**Figure 1.**
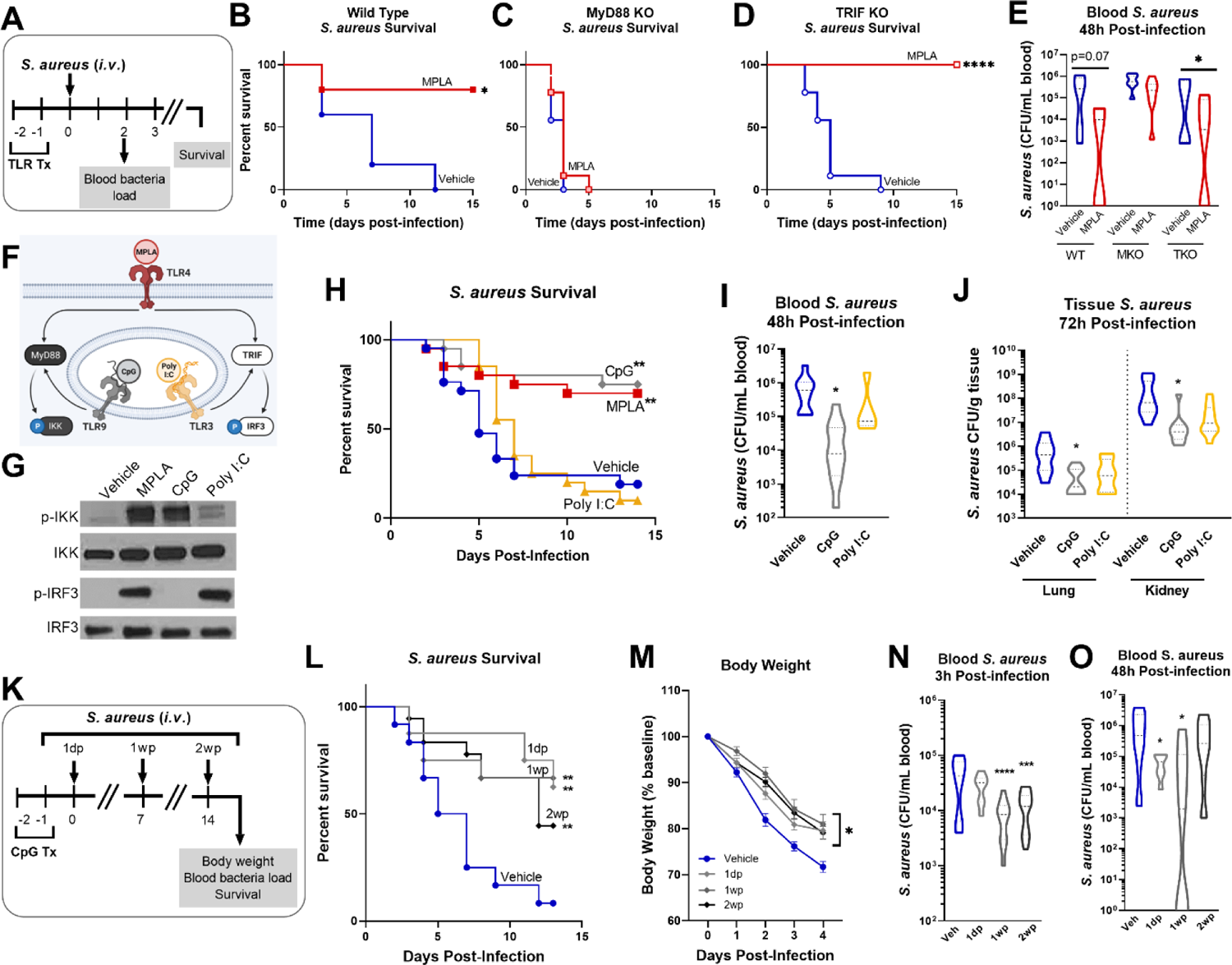
MYD88-activating CpG, but not TRIF-activating Poly I:C, confers resistance to *S. aureus* infection. (**A**) Mice were injected with TLR agonists (20 µg *i.v.*), or vehicle (Lactated Ringers) for two consecutive days prior to systemic challenge with *S. aureus* (*i.v.*). (**B-D**) Wild type (B), MyD88-KO (C), and TRIF-KO mice (D) were challenged with *S. aureus* and survival was monitored for 15 days (n=5-9/group). (**E**) Bacteremia was assessed 48h after infection. **(F**) Schematic of TLR agonist pathway specificity and downstream signaling. (**G**) Pathway specificity of MPLA, CpG, and Poly I:C were determined by Western blotting of protein isolated from wild type BMDMs treated for 24h with TLR agonists compared to unstimulated negative controls. (**H**) Survival was monitored for 15 days post-*S. aureus* inoculation (n = 20-21/group). (**I**) Bacteremia was assessed 48h after infection (n = 5-10). (**J**) Tissue bacterial load (CFU/gram) of lung and kidney was quantified at 72h post-infection (n = 8-10/group). (**K**) Mice were infected with 10^8^ CFU *S. aureus* (*i.v.*) 1 day-post (1dp), 1 week-post (1wp) or 2 weeks-post (2wp) CpG treatment alongside vehicle-treated controls. (**L**) Survival was monitored for 15 days post-infection (n = 10-18/group). (**M**) Body weights were measured at baseline and after infection. (**N-O**) Bacteremia was assessed at 3h (N) and 48h (O) post-infection. Mean ± SEM are shown for body weight (M). * *p* < 0.05, ** *p* < 0.01, **** *p* < 0.0001 by a log-rank Mantel– Cox test for Kaplan–Meier survival plots or otherwise determined by ANOVA followed by Dunnett’s post-hoc multiple comparison test.

Next, we treated mice with MyD88- and TRIF-pathway selective agonists prior to *S. aureus* infection. We chose the TLR9/MyD88-selective agonist CpG-ODN (CpG) and the TLR3/TRIF-selective agonist Poly I:C. As MyD88 activation results in phosphorylation of IkB kinase (p-IKK) and TRIF activation results in phosphorylation of interferon regulatory factor 3 (p-IRF3; **Figure 1F**), we verified pathway specificity by Western blotting of bone marrow derived macrophages (BMDMs) treated with CpG and Poly I:C as compared to the dual-activating agonist MPLA (24h incubation; **Figure 1G**). Mice treated with CpG had significantly higher survival (75% survival) than vehicle controls (20%), a trend similar to MPLA-treated mice (70%); in contrast, Poly I:C-treated mice showed no survival benefit (10%) compared to vehicle controls (**Figure 1H**). CpG treatment reduced bacteremia at 48h post-infection (**Figure 1I**) and bacterial load in the lung and kidneys harvested at 72h post-infection (**Figure 1J**), similarly to MPLA, as previously reported (^13^).

To determine how long CpG-mediated immunoprophylaxis persists, mice were treated with CpG for 2 consecutive days and infected 1 day, 1 week, or 2 weeks-post CpG treatment (1dp, 1wp, and 2wp, respectively; **Figure 1K**). CpG-treated mice had significantly improved survival compared to vehicle-treated controls at 1dp, 1wp, and 2wp-administration (62.5%, 66.7%, and 38.8%, respectively; **Figure 1L**). All CpG-treated groups showed decreased body weight loss in the days following infection compared to controls (**Figure 1M**) and enhanced bacterial clearance at 3h- and 48h-post-inoculation (**Figure 1N, O**). These results demonstrate that MPLA-induced trained immunity is dependent on activation of MyD88, but not TRIF. Further, activation of MyD88-dependent signaling with CpG, but not TRIF-dependent signaling with Poly I:C, confers trained immunity similarly to MPLA and persists for *at least* two weeks after treatment.

### Macrophages drive TLR/MyD88-induced trained immunity and resistance to infection

Our previous work showed that MPLA-induced resistance to infection was not dependent on the adaptive immune system; but was lost after depletion of innate leukocytes (^13^). Since macrophages have a lifespan compatible with the duration of trained immunity, we determined their role in TLR/MyD88-driven trained immunity (^18^). We first depleted macrophages by *i.v.* administration of clodronate-laden liposomes (CL) alongside PBS-laden liposome controls (PL) 24h prior to the first CpG treatment (**Figure 2A**). Macrophage (MΦ)-depleted mice lost the CpG-mediated survival benefit that was observed in macrophage-intact controls. Macrophage depletion also accelerated mortality in Poly I:C-treated mice compared to their macrophage-intact controls (**Figure 2B**). Immunohistochemical staining of F4/80^+^ macrophages in kidneys harvested one-day post-infection demonstrated increased macrophages in CpG-treated, but not Poly I:C-treated, mice compared to vehicle controls in macrophage-intact cohorts. CpG-induced augmentation of kidney macrophages was lost in clodronate-treated mice (**Figure 2C**), demonstrating the efficacy of clodronate in depleting macrophages (low magnification micrographs are provided for further visualization in **Supplemental Figure 1A**). Quantification of F4/80 IHC, conducted by a blinded investigator, statistically validated these observations (**Figure 2C**, right).

**Figure 2.**
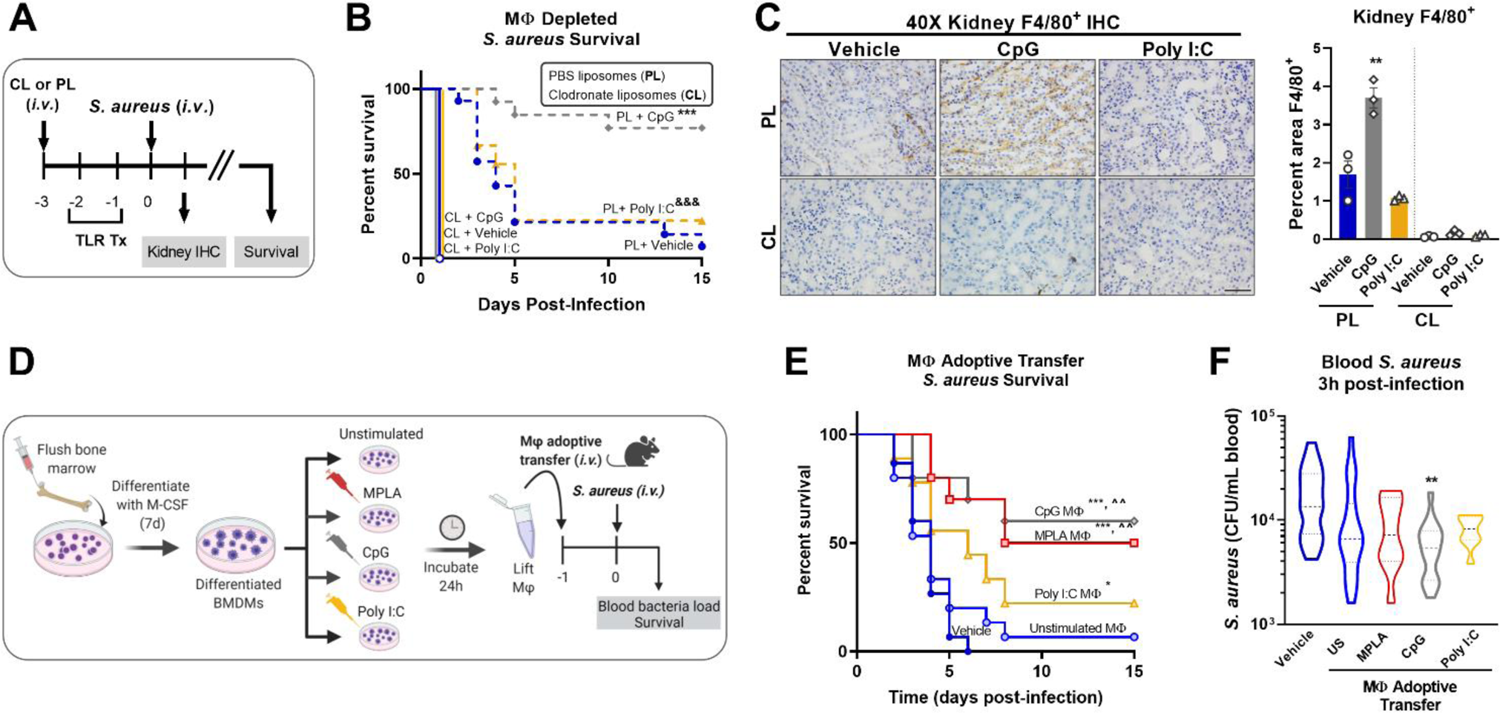
Macrophages are critical for CpG-mediated host resistance to *S. aureus* infection. (**A**) Mice were injected (*i.v.*) with clodronate liposomes (CL) or control PBS liposomes (PL) 24h prior to the first injection of TLR agonist or vehicle. Mice were administered CpG, Poly I:C, or vehicle for two consecutive days (*i.v.*), and challenged with 10^8^ CFU *S. aureus* (*i.v.*) the following day. (**B**) Survival was monitored for 15 days. (**C**) Representative images of F4/80 IHC staining of kidney samples harvested 1-day post-infection of macrophage-depleted (CL) or control (PL) mice are shown (scale bar represents 50 μm); F4/80 IHC was (right, n = 3/group). (**D**) Bone marrow-derived macrophages (BMDMs; MΦ) were treated with MPLA (1 µg/mL), CpG (1 µg/mL), or Poly I:C (10 µg/mL) for 24h or left unstimulated (US), washed, and transferred (5X10^5^ *i.v.*) to otherwise naïve mice 24h prior to systemic challenge with 10^8^ CFU *S. aureus.* (**E**) Survival after *S. aureus* infection in macrophage recipient mice alongside vehicle-treated controls (n=9-15/group) was monitored for 15 days. (**F**) Bacteremia was assessed 3h post-infection. Bars represent mean ± SEM. *** p < 0.001 compared to CL + Vehicle and ^&&&^ p < 0.001 compared to CL + Poly I:C (B), * p < 0.05 and *** p < 0.001 compared to vehicle or ^^compared to unstimulated macrophages (E) by a log-rank Mantel–Cox test for Kaplan–Meier survival plots; ** p < 0.01 compared to vehicle controls determined by ANOVA followed by Dunnett’s post-hoc multiple comparison test.

To further establish the role of macrophages in TLR-mediated infection resistance, BMDMs were treated with MPLA, CpG, or Poly I:C for 24h or left unstimulated as controls, and then adoptively transferred (5X10^5^ *i.v.*) to naïve recipient mice alongside vehicle controls. Mice were infected with *S. aureus* the next day (**Figure 2D**). Mice which received MPLA- or CpG-treated BMDMs had significantly improved survival (50% and 60%, respectively; **Figure 2E**). Poly I:C-treated BMDM-recipient mice showed a slight survival benefit compared to vehicle controls, but were comparable to unstimulated BMDM-recipient controls. CpG-treated BMDMs facilitated significantly improved bacterial clearance (**Figure 2F**) compared to vehicle controls or unstimulated BMDM-recipient controls, with similar but insignificant trends observed for MPLA- and Poly I:C-BMDM-recipient mice. These two approaches demonstrate that TLR/MyD88-mediated protection is driven by macrophages.

### MPLA-induced transcriptional profile is blunted in MyD88- and TRIF-deficient macrophages

To gain further insight into the contributions of MyD88 and TRIF in macrophage trained immunity, we employed an unbiased transcriptomic approach. BMDMs were derived from WT, MyD88-KO and TRIF-KO mice, treated with the dual-activating agonist MPLA or left unstimulated, and RNAseq was performed (**Figure 3A**). Pearson Correlation analysis demonstrates the high consistency of the RNAseq data as indicated by high R^2^ values among biological replicates within groups (**Supplemental Figure 2**). This analysis also shows dissimilarity among MPLA treated samples compared to unstimulated controls. Heat map representation of differentially expressed genes (DEGs) shows a rich transcriptional profile induced by MPLA in WT BMDMs; this was blunted in both MyD88-KO and TRIF-KO macrophages (**Figure 3B**). Still, visual observation of gene clusters shows a similar MPLA-induced expression pattern in WT and TRIF-KO BMDMs, which is divergent in MyD88-KO BMDMs. Principal component (PC) analysis revealed consistency among replicates in each group and clustering among the unstimulated WT, MyD88-KO, and TRIF-KO BMDMs (**Figure 3C**). PC analysis showed unique variance in PC2 in MPLA-treated MyD88-KO BMDMs compared to MPLA-treated WT and TRIF-KO macrophages.

**Figure 3.**
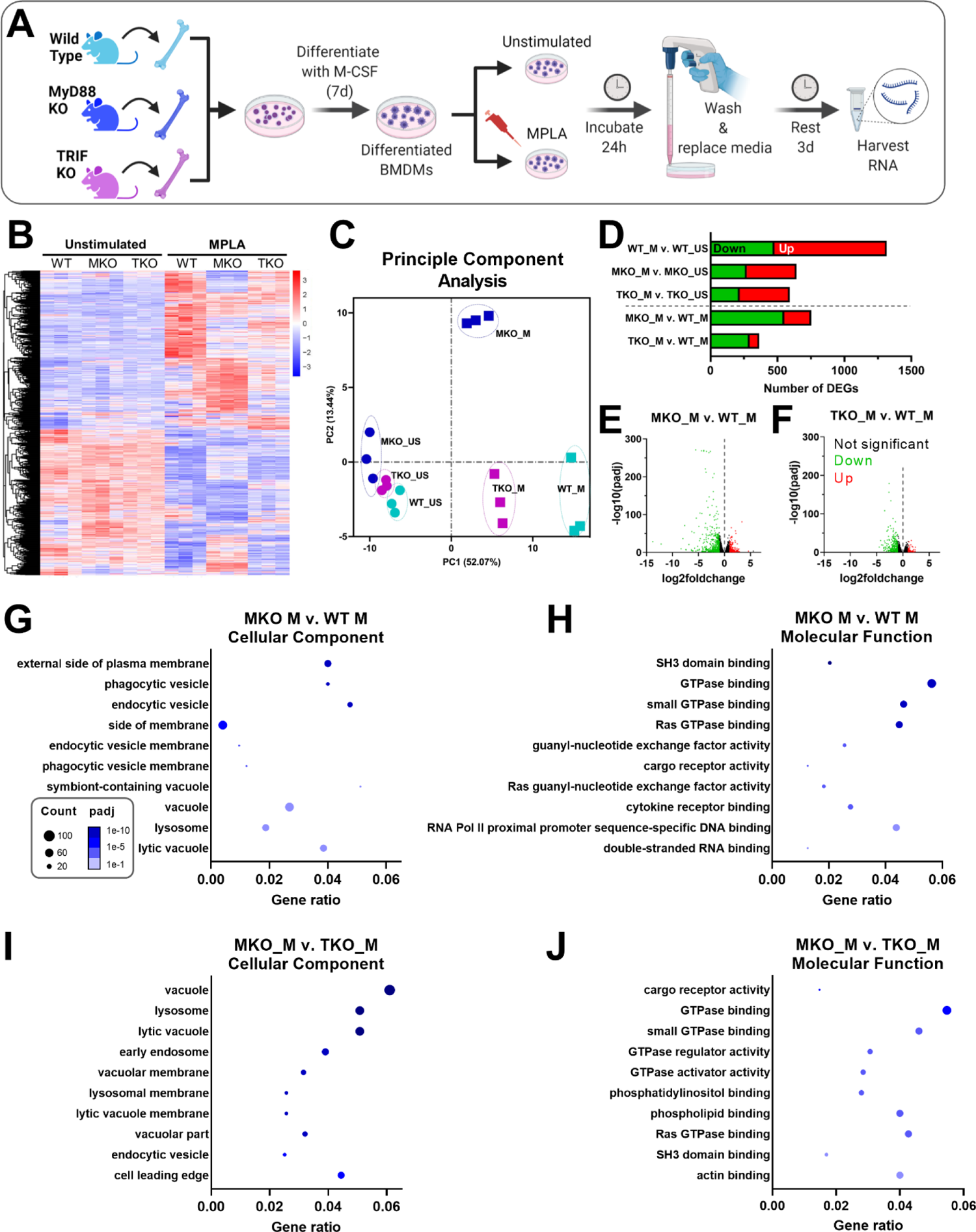
MPLA-induced transcription is blunted in TRIF-KO and MyD88-KO macrophages. (**A**) Bone marrow-derived macrophages (BMDMs) were cultured from WT, MyD88-KO (MKO) and TRIF-KO (TKO) mice. BMDMs were treated with MPLA (1 μg/mL) or left unstimulated (US) as controls. After 24h incubation, media was changed and cells were rested for 3 days before harvesting RNA for RNAseq analysis. (**B**) Differential gene expression analysis of unstimulated control or MPLA-treated WT, MKO, and TKO BMDMs as shown by heatmap of all samples represented by columns (n=3/group) and rows corresponding to different genes. (**C**) Principal component analysis of MPLA-treated (_M) or unstimulated (_US) WT, MKO, and TKO BMDMs was conducted. (**D**) The number of MPLA-induced differentially expressed genes (DEGs) compared to genotype unstimulated controls (above dotted line) or compared to MPLA-treated WT cells (below dotted line) were quantified. (**E-F**) Down-regulated (green) and up-regulated (red) DEGs were visualized by volcano plots for MPLA-treated MKO cells (E) or MPLA-treated TKO cells (F) compared to MPLA-treated WT cells. **G-J** Gene ontology (GO) functional enrichment analysis was conducted for the 10 most downregulated pathways in the cellular component and molecular function classes in MPLA-treated MKO compared to MPLA-treated WT macrophages (G-H), or compared to MPLA-treated TKO macrophages (**I-J**). The number of genes differentially regulated in each respective pathway (*i.e.* “count”) are indicated by dot size and degree of significance (padj) by intensity of color. n=3 biological replicates/group. Log2foldchange >1 and adjusted *p* value <0.05 were considered differentially expressed.

MPLA treatment induced 478 down-regulated and 838 up-regulated genes (total 1,316 DEGs) in WT cells, which was blunted in both MyD88-KO (total 643 DEGs) and TRIF-KO macrophages (total 592 DEGs; **Figure 3D**). The degree of transcriptional similarity among MPLA-treated KO BMDMs compared to MPLA-treated WT BMDMs (Figure 1D below dashed line) was assessed. When compared to MPLA-treated WT cells, MPLA-treated MyD88-KO BMDMs had 752 DEGs, whereas MPLA-treated TRIF-KO macrophages had only 352 DEGs, suggesting that MPLA-induced transcription is more conserved in TRIF-KO macrophages than in MyD88-KO macrophages. Volcano plots were used to visualize the magnitude of DEGs in MPLA-treated KO vs MPLA-treated WT BMDMs. These scatterplots show that the DEGs in MPLA-treated MyD88-KO macrophages were greater in magnitude and were more significant than the DEGs in MPLA-treated TRIF-KO BMDMs compared to MPLA-treated WT BMDMs (**Figure 3E** and **3F**, respectively).

To elucidate which transcriptomic changes are essential in MPLA-induced trained immunity, we performed enrichment analyses. Since MPLA-induced training is lost in MyD88-KO macrophages, we queried the most downregulated enrichment pathways in MPLA-treated MyD88-KO BMDMs compared to MPLA-treated WT cells. We identified the 5 most downregulated enrichment pathways in each gene ontology (GO) class (biological process, cellular component, and molecular function; **Supplemental Figure 3**). We also evaluated these pathways in MPLA-treated TRIF-KO BMDMs since MPLA-induced trained immunity is functionally preserved in TRIF-KO cells. Downregulation of GO pathways in the biological process class were similarly altered in both MPLA-treated MyD88-KO and TRIF-KO cells. Conversely, MPLA-treated TRIF-KO BMDMs showed less deviation from MPLA-treated WT cells than MPLA-treated MyD88-KO BMDMs in pathways belonging to the cellular component and molecular function classes, indicating that these processes are most critical for MPLA-induced trained immunity.

To further investigate these enrichment pathways, we generated dot plot representations of the 10 most downregulated pathways in the cellular component or molecular function classes when comparing MPLA-treated MyD88-KO macrophages to either MPLA-treated WT or MPLA-treated TRIF-KO macrophages. Dot plots show gene ratios (number of DEGs in a given pathway relative to total genes) on the x-axis, number of genes in that pathway that are differentially regulated by dot size, and degree of pathway significance by dot color. The most downregulated pathways in the cellular component class in MPLA treated MyD88-KO cells included phagocytic vesicle and lysosome (**Figure 3G**), and the most downregulated molecular function pathways were related to GTPase binding and activity (**Figure 3H**). These were similarly downregulated in MPLA treated MyD88-KO cells compared to MPLA-treated TRIF-KO macrophages (**Figure 3I-J**), indicating that these pathways are critical for MPLA-induced trained immunity.

### TLR-induced macrophage metabolic reprogramming is dependent on MyD88

As demonstrated by our group and others, sustained metabolic rewiring is a key characteristic of trained immunity (^5, 12, 13, 16, 19^). We previously showed elevation of glycolysis and mitochondrial respiration were similar in CpG-treated and MPLA-treated BMDMs relative to unstimulated controls (^12^) suggesting that MyD88 signaling plays a role in TLR-mediated metabolic augmentation. We assessed metabolic function after stimulation with MyD88- and TRIF-stimulating ligands. Here, BMDMs were treated with TLR agonists for 24h and assessed immediately (24h; acute changes) or agonists were washed off after the 24h incubation and assessed at 3 days post-treatment (3dp; indicative of sustained changes; **Figure 4A**). Previous studies show that acute exposure to TLR4 agonists triggers aerobic glycolysis (*i.e.* the Warburg effect) whereby glycolysis is elevated without a change in oxygen consumption; while trained macrophages have sustained increased glycolysis which is accompanied by increased mitochondrial respiration (^13^). We aimed to determine the roles of MyD88- and TRIF-dependent signaling in this phenotype. Twenty-four-hour incubation of WT BMDMs with MPLA or CpG prompted a sharp increase in the extracellular acidification rate (ECAR, indicative of glycolytic capacity) which was further elevated in 3dp groups (approximately 10-fold higher than unstimulated controls), whereas the ECAR of Poly I:C-treated BMDMs were comparable to controls (**Figure 4B**). Assessment of mitochondrial respiration measured by oxygen consumption rate (OCR) showed similar trends (**Figure 4C**).

**Figure 4.**
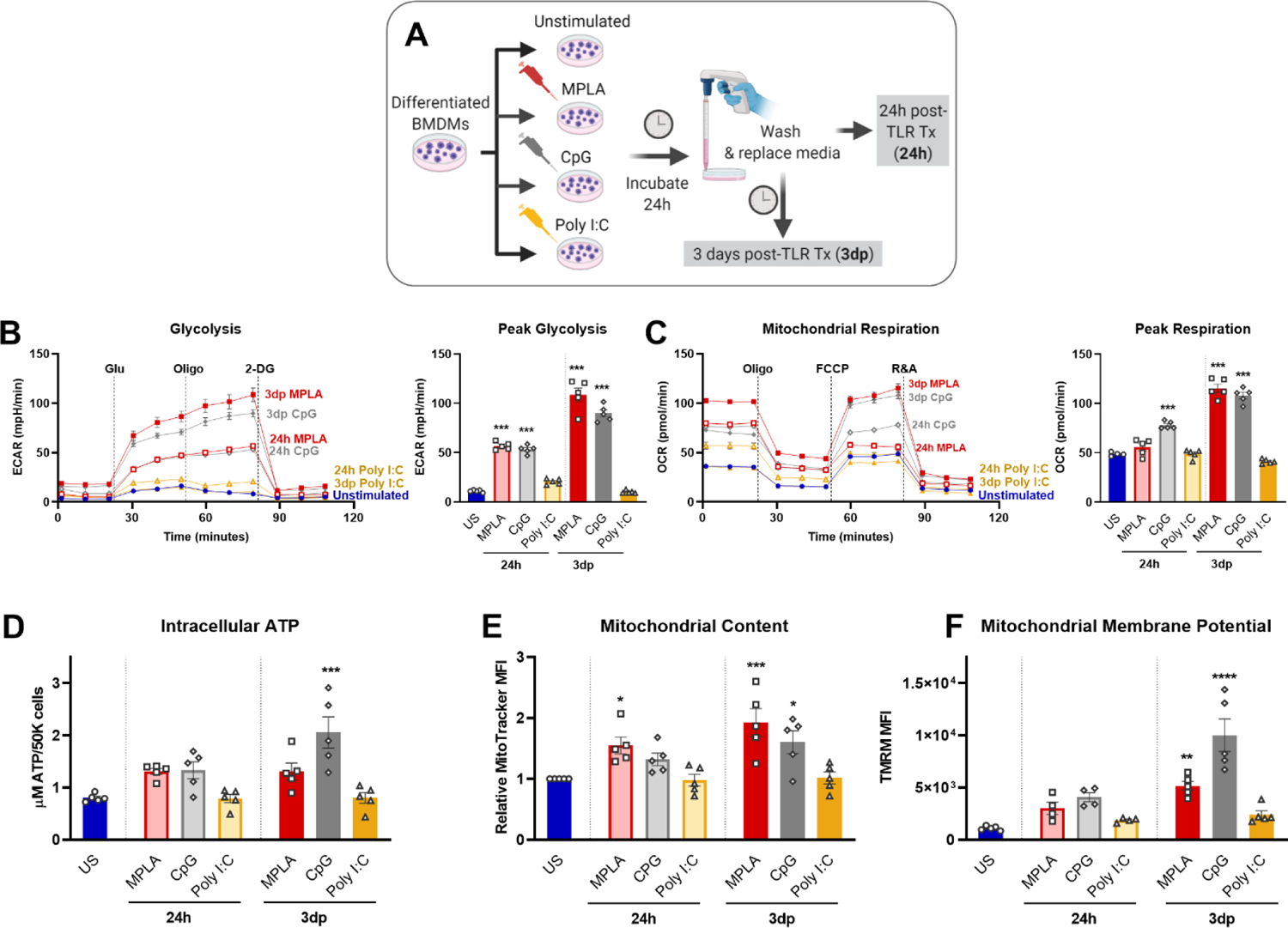
MyD88-selective CpG, but not TRIF-selective Poly I:C, triggers macrophage metabolic reprogramming. (**A**) Differentiated BMDMs were treated with MPLA (1 µg/mL), CpG (1 µg/mL), Poly I:C (10 µg/mL) or left unstimulated as controls for 24h. BMDMs were either washed and immediately assayed (24h) or rested for 3 days prior to assay (3dp). (**B**) Glycolysis stress test was performed and extracellular acidification rate (ECAR) was measured by Seahorse Xf^e^96 (left); peak glycolysis was analyzed (right). (**C**) Mitochondrial stress test of BMDMs was performed and oxygen consumption rate (OCR) was measured (left); peak respiration was analyzed (right). (**D**) BMDMs (5X10^4^) were washed, lysed, and ATP content was measured. (**E**) Mitochondrial content was assessed using MitoTracker Green staining and (**F**) mitochondrial membrane potential was assessed using TMRM staining. Data are shown as mean ± SEM, n = 4-5/group * *p* < 0.05, ** *p* < 0.01, *** *p* < 0.001, **** *p* < 0.0001 by ANOVA followed by Dunnett’s post-hoc multiple comparison test.

Further, we found that 3dp MPLA- and CpG-treated BMDMs showed trends of increased intracellular ATP compared to unstimulated controls or Poly I:C-treated cells (**Figure 4D**). MPLA- and CpG-treated, but not Poly I:C-treated BMDMs showed increased mitochondrial content (**Figure 4E**) indicating mitochondrial biogenesis, and robust elevation of mitochondrial membrane potential, especially in the 3dp groups (**Figure 4F**). We verified that activation of MyD88 via TLR2 induced similar trends in intracellular ATP, mitochondrial content, and mitochondrial activity using the TLR2/MyD88-specific agonist Pam3CSK (**Supplemental Figure 4A-C**).

We next assessed the relative contribution of MyD88 and TRIF signaling in driving metabolic rewiring on a transcriptomic and functional level. RNAseq analysis showed that metabolism-related GO pathways, such as pyruvate metabolism and citrate cycle, were similarly induced by MPLA in WT and TRIF-KO BMDMs, but not MyD88-KO BMDMs (**Figure 5A**). Expression of genes in these metabolism-related enrichment pathways (*e.g.* electron transport activity, glycolytic process, and regulation of glucose transport) showed similar induction in MPLA-treated WT and TRIF-KO BMDMs, but was blunted in MyD88-KO BMDMs (**Figure 5B**). Hierarchal clustering analysis of these metabolism-related genes demonstrated further distance of MPLA-treated MyD88-KO BMDMs compared to MPLA-treated WT and TRIF-KO BMDMs (**Supplemental Figure 5**). Functionally, we found that dual-activating MPLA and MyD88-activating CpG triggered increased ECAR in TRIF-KO BMDMs similarly to that in WT, which was not observed in MyD88-KO cells (**Figure 5C-D**). Similar trends were observed for mitochondrial respiration (**Figure 5E-F**). We confirmed that the MyD88-stimulating TLR2 agonist Pam3CSK increased glycolysis and mitochondrial respiration in TRIF-KO cells similarly to WT, a trend which was absent in MyD88-KO BMDMs (**Supplemental Figure 4D-E**). Together, these data demonstrate that MyD88, but not TRIF, is critical for TLR-induced macrophage metabolic rewiring.

**Figure 5.**
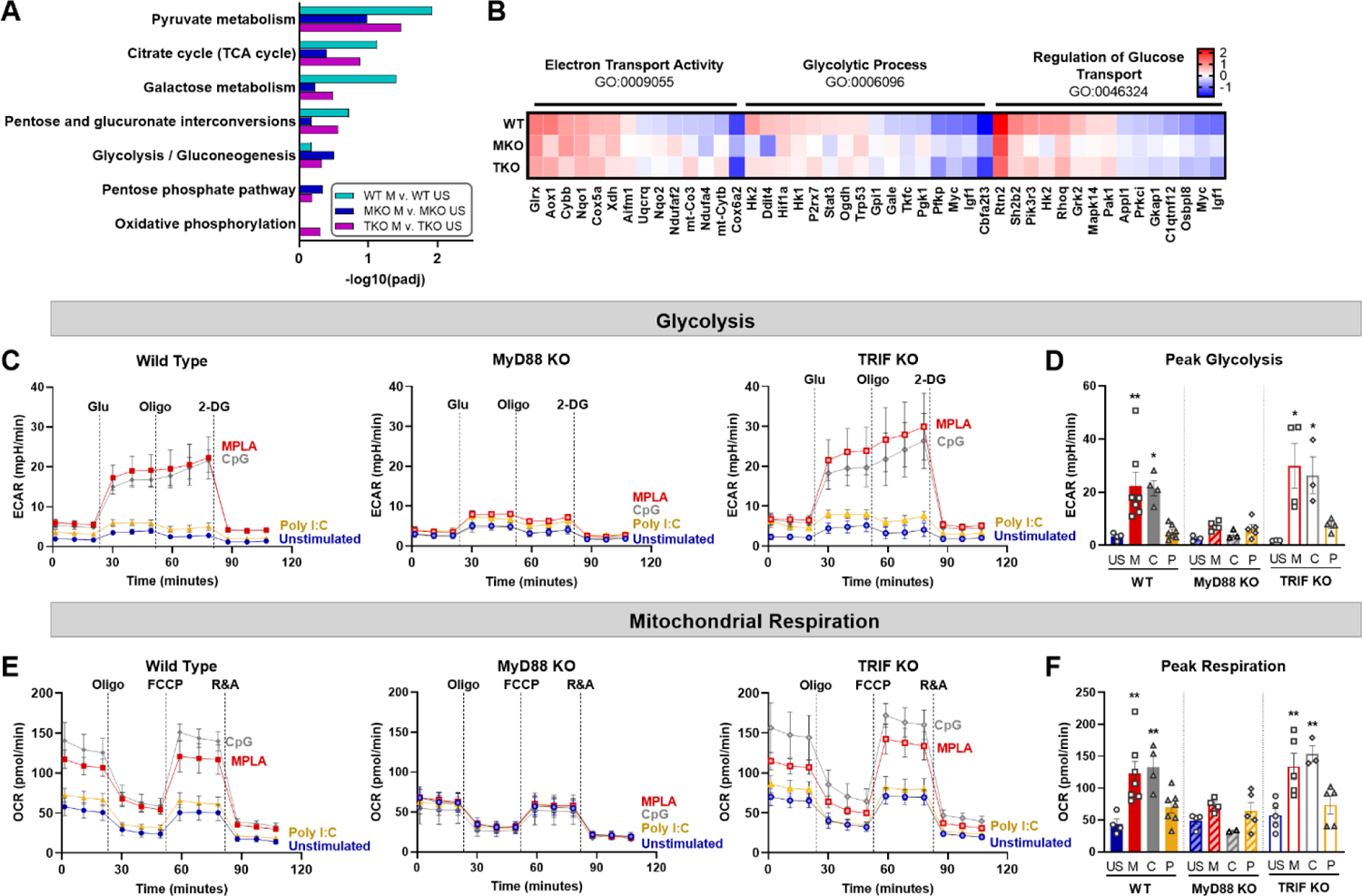
TLR-induced metabolic rewiring is lost in MyD88-deficient macrophages. (**A-B**) Gene Ontology (GO) functional enrichment analysis was conducted on MPLA-treated WT, MyD88-KO (MKO), and TRIF-KO (TKO) bone marrow-derived macrophages (BMDMs) for metabolism-related pathways and displayed by significance of pathway induction (A). Differential expression of genes in metabolism-related GO pathways are shown by heatmap (B). (**C-D**) Glycolysis stress test was performed using Seahorse Xf^e^96 which measured extracellular acidification rate (ECAR) of TLR agonist-treated (MPLA, M; CpG, C; Poly I:C, P) or unstimulated (US) WT (left) MKO (middle), and TKO (right) BMDMs (C). Peak glycolysis was measured (D). (**E-F**) Mitochondrial stress test was performed under the same experimental parameters and oxygen consumption rate (OCR) is presented for WT (left), MKO (middle), and TKO (right) BMDMs (E). Peak respiration was measured (F). Data are shown as mean ± SEM, n = 3-5/group * *p* < 0.05, ** *p* < 0.01 by ANOVA followed by Dunnett’s post-hoc multiple comparison test.

### TLR-induced augmentation of macrophage antimicrobial function is mediated by MyD88 signaling

We performed enrichment analyses on MPLA-treated WT, MyD88-KO, and TRIF-KO macrophages to determine the role of these signaling cascades in TLR-mediated augmentation of antimicrobial functions. We found that pathways such as leukocyte differentiation, positive regulation of cytokine production, and leukocyte proliferation were similarly enriched by MPLA treatment in WT and TRIF-KO BMDMs, but were less enriched in MPLA-treated MyD88-KO cells (**Figure 6A**). A dot plot was also generated which showed that gene expression ratios were consistently lower in MPLA-treated MyD88-KO BMDMs compared to WT and TRIF-KO BMDMs (**Figure 6B**). We found that MPLA-induced transcriptional regulation of phagocytosis, ROS metabolic process, and chemotaxis were largely similar among TRIF-KO and WT macrophages, but MPLA-induction of these terms were largely blunted in MyD88-KO cells (**Figure 6C**). To further understand the correlation among these groups, clustering analysis of the listed antimicrobial-related genes was performed which demonstrated further distance of MPLA-treated MyD88-KO BMDMs compared to MPLA-treated WT and TRIF-KO BMDMs (**Supplemental Figure 6**). qRT-PCR confirmed that CpG induces antimicrobial-related transcription similarly to MPLA as *MARCO* (**Figure 6D**), *PECAM1* (**Figure 6E**), and *Ptprm* (**Figure 6F**) expression were upregulated by MPLA and CpG in WT and TRIF-KO BMDMs. These trends were largely absent in MyD88-KO BMDMs with the exception of modest upregulation of *MARCO*. Treatment with Poly I:C had no observed effect on transcription levels of these genes.

**Figure 6.**
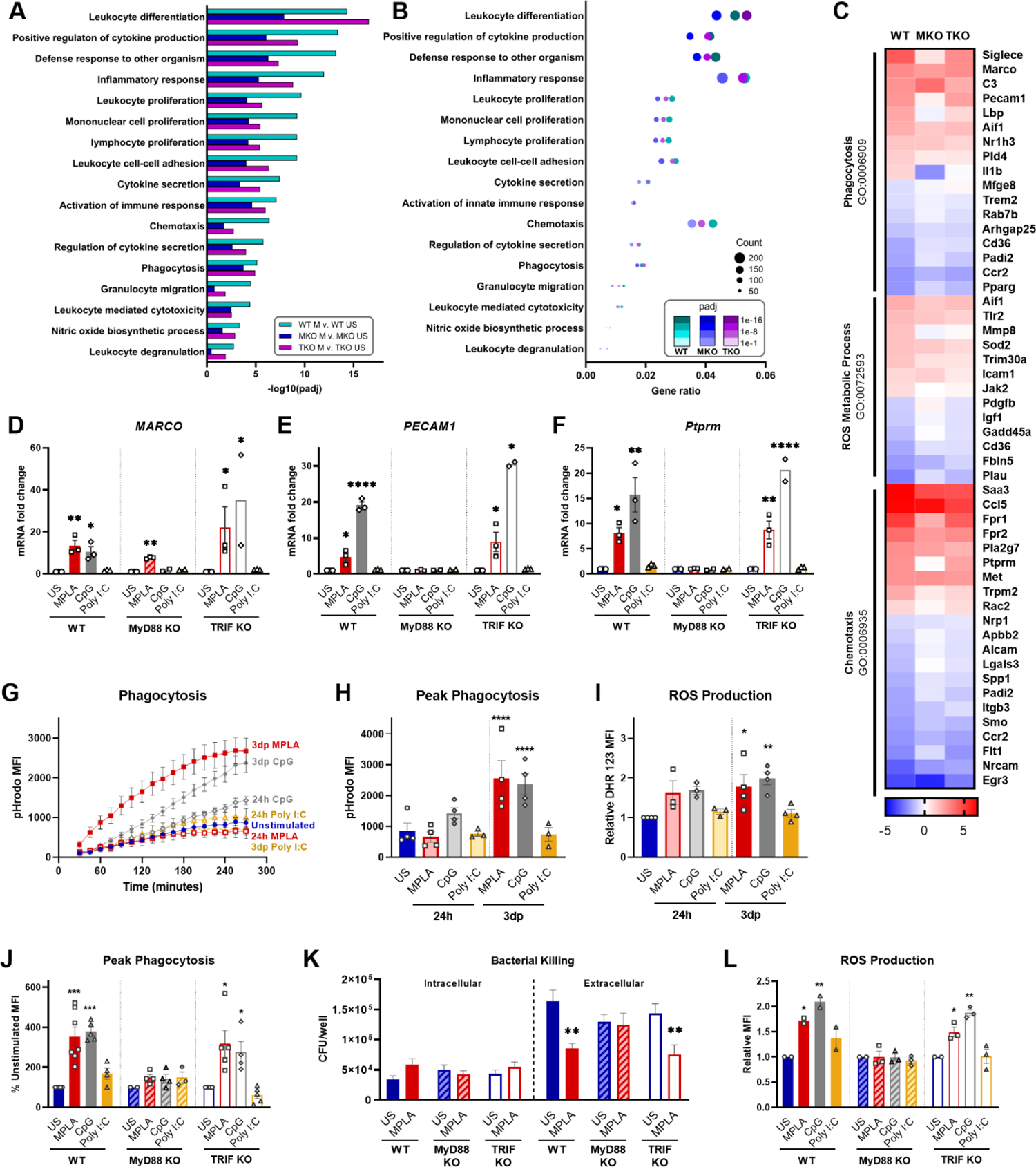
TLR agonist-induced augmentation of antimicrobial capacity is blocked in MyD88-KO, but not TRIF-KO, macrophages. (**A-B**) Gene ontology (GO) functional enrichment analysis was conducted for immunology-related pathways and displayed by significance of pathway induction (A) and by dot plot which shows number of genes differentially regulated in each respective pathway (*i.e.* “count”) by dot size and degree of significance (padj) by intensity of color (B). (**C**) Immune function-related GO pathways were evaluated by heat map comparing MPLA-trained WT, MyD88-KO (MKO), and TRIF-KO (TKO) BMDMs to unstimulated (US) genotype controls. (**D-F**) Relative gene expression of *MARCO* (D), *PECAM1* (E) and *Ptprm* (F) of unstimulated control, MPLA, CpG, and Poly I:C-treated WT, MKO, and TKO BMDMs 3 days post-treatment (n=2-3 biological replicates/group). (**G-H**) The fluorescence emitted upon phagocytosis of pHrodo-tagged *S. aureus* bioparticles was measured over 4 hours (G) and peak phagocytosis was analyzed (H). (**I**) ROS production was assessed by flow cytometry. (**J**) Phagocytic capacity of MPLA-, CpG-, and Poly I:C-treated WT, MyD88-KO, and TRIF-KO were assessed alongside unstimulated (US) genotype controls. (**K**) Bacterial killing (*S. aureus* MOI 10) of MPLA-treated WT, MyD88-KO, and TRIF-KO BMDMs compared to US controls. (**L**) ROS production capacity was assessed. Data are shown as mean ± SEM, * *p* < 0.05, ** *p* < 0.01, *** *p* < 0.001, *** *p* < 0.0001 by ANOVA followed by Dunnett’s post-hoc multiple comparison test.

Next, we evaluated the role MyD88 and TRIF in TLR-augmentation of antimicrobial functions. Phagocytic capacity was assessed using pHrodo-tagged *S. aureus* bioparticles which emit fluorescence upon internalization into the phagolysosome; this assay showed significantly increased phagocytosis (3-fold higher at peak activity) by 3dp MPLA- and CpG-, but not Poly I:C-treated macrophages (**Figure 6G, H**). Production of antimicrobial ROS followed a similar trend (**Figure 6I**).

We further assessed antimicrobial function of MyD88-KO and TRIF-KO BMDMs compared with WT controls at 3 days post-treatment with the respective agonists. We observed increased phagocytosis by WT and TRIF-KO BMDMs after MPLA or CpG treatment compared to unstimulated controls, an effect that was lost in MyD88-KO BMDMs (**Figure 6J**). Complimentary to this finding, in vitro *S. aureus* killing assay showed that MPLA-treated WT and TRIF-KO BMDMs had unchanged intracellular bacteria but significantly reduced extracellular bacteria, indicative of increased bacterial killing, which was not observed in MyD88-KO BMDMs (**Figure 6K**). MPLA and CpG treatment also increased ROS production compared to unstimulated controls (**Figure 6L**). No change in these antimicrobial functions was observed by treatment with Poly I:C or in BMDMs deficient in MyD88. Activation of TLR2, which is a MyD88-selective cell surface receptor, using Pam3CSK resulted in similar trends (**Supplemental Figure 7**). Taken together, these data indicate that MyD88, but not TRIF, is critical for TLR-induced augmentation of antimicrobial functions in macrophages.

### mTOR activation is required, in part, for TLR/MyD88-induced macrophage reprogramming

Previously, we provided evidence that blockade of mTOR signaling prevented MPLA-mediated macrophage reprogramming as well as MPLA-induced infection resistance (^13^). Since mTOR is a key regulator of cellular metabolism, we sought to determine if mTOR is also required for CpG-mediated macrophage reprogramming. We first evaluated whether MyD88-stimulating CpG and TRIF-stimulating Poly I:C activate mTOR by measuring S6K phosphorylation. p-S6K was significantly increased in CpG-treated BMDMs (2.1-fold change) similarly to MPLA-treated cells (2.8-fold change); however, p-S6 was not significantly changed in Poly I:C-treated macrophages compared to unstimulated controls (1.2-fold change; **Figure 7A**).

**Figure 7.**
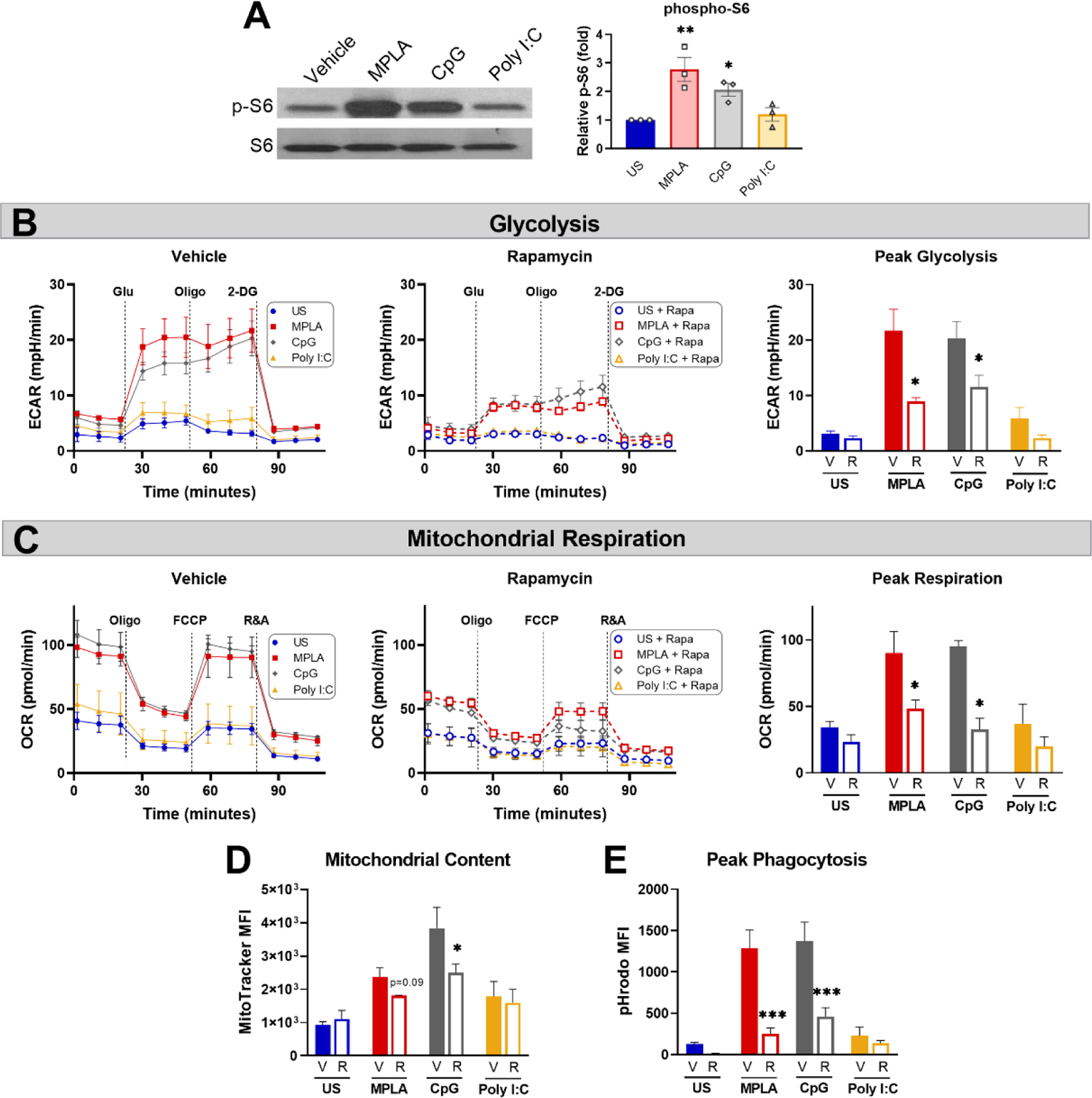
mTOR is a key mediator of TLR/MyD88-induced macrophage metabolic and functional reprogramming. (**A**) Western blotting of phosphorylated S6 kinase (p-S6) and total S6K (S6) on protein lysates from BMDMs treated for 24h with the TLR agonists (1 µg/mL for MPLA and CpG, 10 µg/mL for Poly I:C) compared to unstimulated negative controls (left). Blots are representative of three repeated experiments which were quantified by densitometry analysis (right). (**B-E**) BMDMs were treated with 100 nM Rapamycin (R) or vehicle (V) 1h prior to the addition of TLR agonists which remained in culture until washed off 24h later and cells were rested for 3 days prior to assays. (**B**) Glycolysis stress test was performed using Seahorse Xfe96 which measured extracellular acidification rate (ECAR) for vehicle-treated (left) and rapamycin-treated (middle) BMDMs, and peak glycolysis was analyzed (right). (**C**) Mitochondrial stress test was performed under the same experimental parameters as glycolysis stress test, and oxygen consumption rate (OCR) is presented for vehicle-treated (left), rapamycin-treated (middle), and peak respiration was measured (right). (**D**) Mitochondrial content was assessed using MitoTracker Green staining. (**E**) Phagocytic capacity was indicated by fluorescence emitted upon phagocytosis of pHrodo-tagged *S. aureus* bioparticles and analyzed at peak activity. Data are shown as mean ± SEM, n = 2-3/group * *p* < 0.05, ** *p* < 0.01, *** *p* < 0.001 by ANOVA followed by Dunnett’s post-hoc multiple comparison test.

To study the effect mTOR inhibition on TLR/MyD88-induced macrophage reprogramming, BMDMs were treated with rapamycin or vehicle 1h prior to TLR agonist treatment. Rapamycin remained in culture media for the 24h incubation after which cells were washed and rested 3 days prior to metabolic and antimicrobial function assays. Rapamycin significantly attenuated MPLA- and CpG-induced augmentation of glycolysis (**Figure 7B**) and mitochondrial respiration (**Figure 7C**) compared to vehicle + TLR agonist-treated controls. Likewise, rapamycin blunted TLR/MyD88-mediated increased mitochondrial content (**Figure 7D**). Additionally, rapamycin prevented MPLA- and CpG-mediated increased phagocytic capacity (**Figure 7E**). Together, these data indicate that dual-pathway activating MPLA and MyD88-stimulating CpG, but not TRIF-stimulating Poly I:C, activate mTOR to facilitate TLR-mediated macrophage reprogramming.

### CpG triggers macrophage metabolic and functional rewiring in vivo

Our next goal was to determine whether TLR-induced functional programs observed in vitro are induced in vivo and to assess how long alterations persist. Mice were treated with CpG for two consecutive days (20 µg *i.v.*) and euthanized 1 day (1d), 3 days (3d), 1 week (1w), or 2 weeks (2w) post-treatment at which time spleens were harvested, weighed, and processed for magnetic isolation of F4/80^+^ macrophages (**Figure 8A**). We observed splenomegaly at 1-week post-CpG treatment, which was resolved by 2 weeks (**Supplemental Figure 8A**) which correlated with increased numbers of isolated macrophages (data not shown). Magnetic isolation of F4/80^+^ macrophages resulted in an enriched population (>85%, **Supplemental Figure 8B**) with high viability and yield for downstream assays. MitoTracker Green staining revealed that mitochondrial content is increased in tissue resident macrophages for one week after in vivo treatment with CpG, indicating mitochondrial biogenesis (**Figure 8B**). A similar trend was observed for intracellular ATP content (**Figure 8C**). Phagocytic capacity was assessed by incubation of isolated splenic macrophages with pHrodo-tagged *S. aureus* bioparticles which showed increased MFI for 1-week post-CpG treatment (**Figure 8D**). Capacity of ROS production was also measured by flow cytometry, which was increased for one week, but was comparable to vehicle controls by two weeks post-CpG treatment (**Figure 8E**). These data demonstrate that the metabolic and antimicrobial phenotype well-described using in vitro models is similarly triggered in vivo. Further, such rewiring occurs for at least one week following administration.

**Figure 8.**
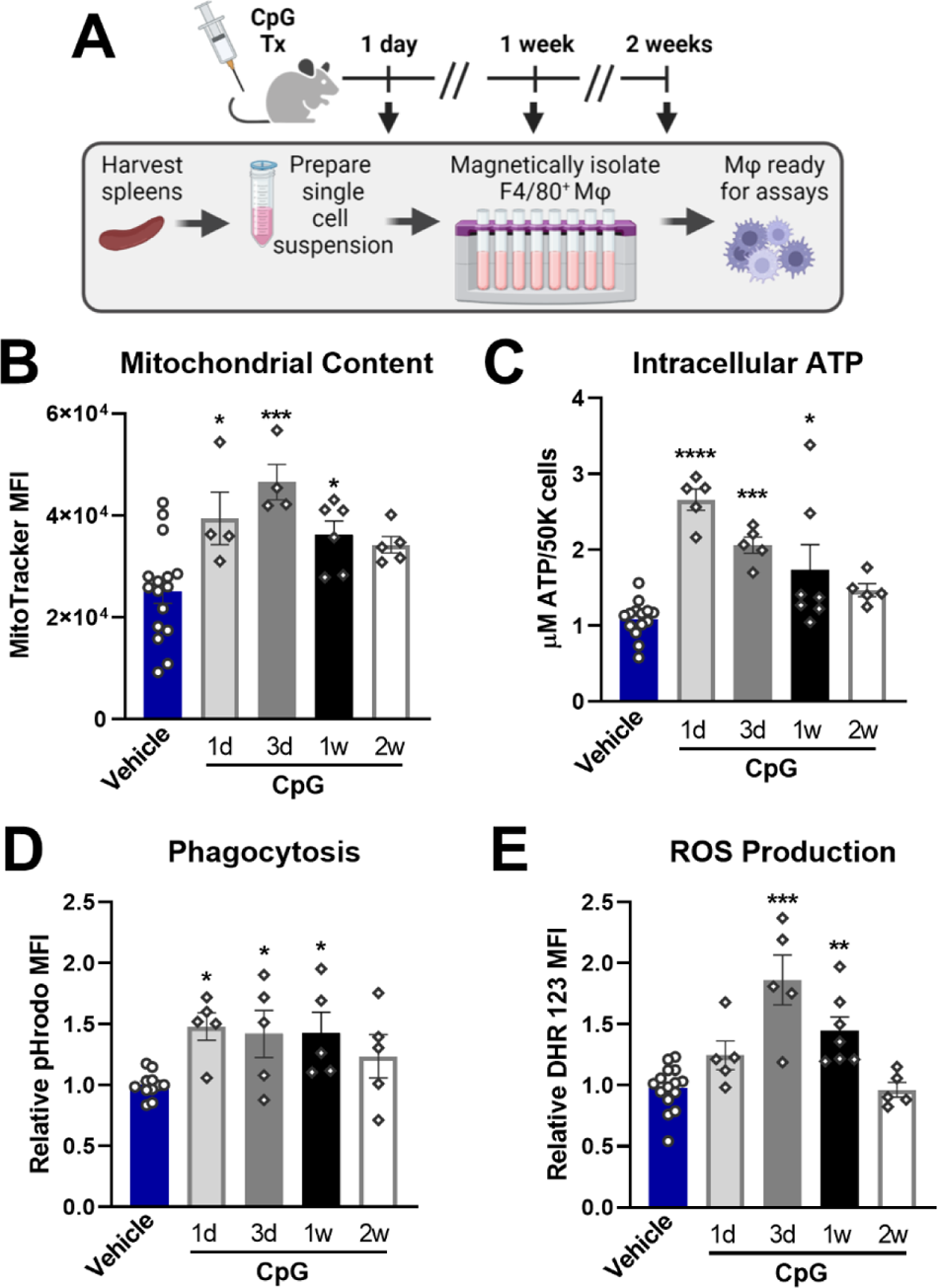
CpG induces macrophage metabolic and functional rewiring *in vivo*. (**A**) Mice were treated for two consecutive days with CpG (20 µg *i.v.*) or vehicle (Lactated Ringers) and spleens were harvested 1-day (1d), 3-days (3d), 1-week (1w), or 2-weeks (2w) later. Spleens were processed for magnetic isolation of F4/80^+^ macrophages which were used immediately for respective assays. (**B**) Mitochondrial content was assessed by MitoTracker Green staining. (**C**) Intracellular ATP content was measured. (**D**) Relative phagocytic activity was assessed by emitted fluorescence upon phagocytosis of pHrodo-tagged *S. aureus* bioparticles and measured by flow cytometry. (**E**) ROS production was determined by relative dihydrorhodamine 123 (DHR 123) fluorescence. Data are shown as mean ± SEM, n = 5-16/group * *p* < 0.05, ** *p* < 0.01, *** *p* < 0.001, **** *p* < 0.0001 by ANOVA followed by Dunnett’s post-hoc multiple comparison test.

### TLR/MyD88 activation induces trained immunity in human monocyte-derived macrophages

To ascertain the translational impact of our studies, we determined whether TLR agonists induce reprogramming via activation of MyD88 signaling in human monocyte-derived macrophages. We obtained buffy coat samples from healthy males from which CD14^+^ monocytes were magnetically isolated and differentiated into macrophages. The resulting human monocyte-derived macrophages (hMDMs) were treated with TLR agonists according to the same timeline as BMDM studies (treated for 24h, washed off, and rested for 3 days prior to assay). We found that dual-activating MPLA and MyD88-activating CpG, but not TRIF-activating Poly I:C, increased glycolytic metabolism (**Figure 9B**),quantified as peak glycolysis (**Figure 9C**). Likewise, MPLA and CpG treatment, but not Poly I:C treatment, resulted in increased mitochondrial respiration (**Figure 9D**),quantified as maximal respiration (**Figure 9E**), and mitochondrial biogenesis evidenced by increased mitochondrial content (**Figure 9F**). Similar trends were observed for the antimicrobial functions of phagocytosis (**Figure 9G**) and ROS production (**Figure 9H**). These data indicate that TLR-mediated macrophage reprogramming occurs in human-derived samples, and that MyD88-, but not TRIF-activation, is required.

**Figure 9.**
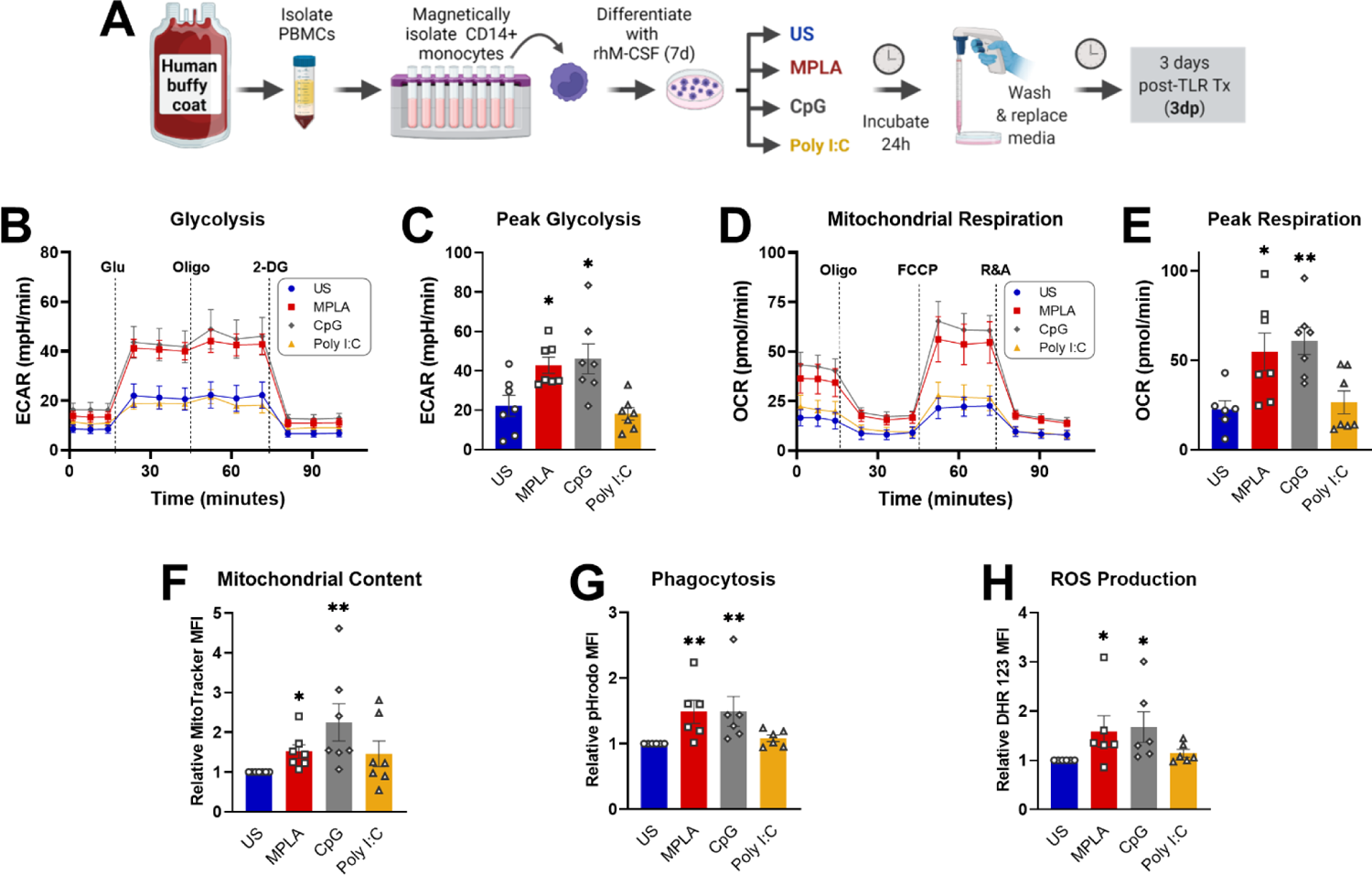
Activation of TLR/MyD88 signaling induces training in human monocyte-derived macrophages. (**A**) PBMCs were enriched from human buffy coat samples and monocytes were isolated using immunomagnetic CD14^+^ positive selection. Monocytes were differentiated for 7d with recombinant human M-CSF (rhM-CSF) resulting in human monocyte-derived macrophages (hMDMs). hMDMs were treated for 24h with MPLA (10 µg/mL), CpG (10 µg/mL), or Poly I:C (100 µg/mL) or left unstimulated (US) as negative controls. Agonist-treated hMDMs were washed, media was replaced, and cells were rested for 3 days prior to assays (i.e. 3 days post-treatment; 3dp). (**B**) Extracellular acidification rate (ECAR) was measured as an indicator of glycolytic capacity and (**C**) Peak glycolysis was analyzed. (**D**) Oxygen Consumption Rate (OCR) was measured and (**E**) peak mitochondrial respiration were analyzed. (**F**) Mitochondrial content was assessed by MitoTracker staining. (**G**) Phagocytic capacity was measured by fluorescence of pHrodo-labeled *S. aureus*. (**H**) ROS production was determined by relative dihydrorhodamine 123 (DHR 123) fluorescence. N=6-7/group, * *p* < 0.05 and ** *p* < 0.01 by ANOVA followed by Dunnett’s post-hoc multiple comparison test.

## DISCUSSION

Investigation of trained immunity has been an active field of research with major discoveries advancing the field towards therapeutic translation for several applications; however, much remains to be explored before the true immunomodulatory potential of training agents can come to fruition. Toll-like receptor (TLR)-4 agonists have been shown to trigger host resistance to infection that lasts for up to two weeks in several clinically relevant models of infection (^10, 11, 15^), which our group has revealed to be driven by long-term changes in innate immune function supported by metabolic rewiring (^13^). We sought to define which signaling pathway(s) mediate immune training given that TLR4 uniquely signals through both MyD88- and TRIF-dependent signaling cascades. Previous research found that TRIF-, rather than MyD88-dependent signaling, is critical for MPLA to mediate augmentation of vaccine efficacy as an adjuvant (^17^). On the other hand, our previous evidence suggested that MyD88 stimulation with CpG, but not TRIF stimulation with Poly I:C, induced protection against an acute intraperitoneal infection with the Gram-negative *P. aeruginosa* (^12^). This prompted us to conduct further investigation regarding these pathways in TLR-mediated trained immunity. Elucidating responsible signaling mechanisms could potentially identify therapeutic targets downstream of TLR activation to circumvent the potential concern regarding unwanted activation of inflammatory and/or autoimmune dysfunction by application of TLR ligands as drugs.

In the current study, we provide evidence that MyD88-selective, but not TRIF-selective, TLR agonists confer protection against *S. aureus* infection for at least two weeks in a macrophage-dependent manner. This correlates with our previous reports that TLR4 agonists can also mediate protection against acute Gram-negative infection for up to two weeks (^12, 15^). Previous data from our laboratory led us to the conclusion that innate leukocytes, rather than T and B lymphocytes, are essential for MPLA-mediated protection against *P. aeruginosa* or *S. aureus* infection (^12, 13^). Numerous trained immunity studies focus on monocytes, however our work previously showed that MPLA-induced resistance to infection is preserved in CCR2 knockout mice which have defective monocyte recruitment. In contrast, protection was lost in mice depleted of macrophages (^13^). Further, our lab has demonstrated that macrophages are essential for mediating trained immunity by β-glucan, a classic inducer of trained immunity (^20^). Together, these results suggest that tissue resident macrophages are critical contributors to TLR-mediated infection resistance. Thus, we hypothesized that macrophages are essential to protection conferred by TLR/MyD88-stimulating agonists. We found that macrophage depletion by clodronate-laden liposomes rendered mice completely vulnerable to infection despite CpG treatment. Complementary to this finding, our data show that adoptive transfer of MPLA- and CpG-trained macrophages, but not Poly I:C-treated macrophages, to naïve mice is sufficient to induce infection prophylaxis. Together, these two approaches establish the clear role of macrophages in driving TLR/MyD88-mediated protection.

It would be of interest to elucidate how long MPLA and CpG can confer infection resistance beyond the 2-week window, especially with growing evidence that training agents induce epigenetic modifications in myeloid progenitors which may propagate host protection long-term (^21^). *de Laval et. al.* have demonstrated that the classic TLR4 ligand lipopolysaccharide (LPS) induces persistent alterations in accessibility of myeloid lineage enhancers in hematopoietic stem cells which is associated with long-term protection against *P. aeruginosa* infection (^22^). We would postulate that the LPS-derivative MPLA and MyD88-activating agonist CpG can induce similar alterations to the epigenetic landscape of myeloid progenitors and long-term infection resistance, which warrants future investigation. Additionally, it would be of interest to test the efficacy of CpG in conferring infection resistance in other clinically relevant models. Based on our prior studies in our 2-hit burn wound infection model whereby MPLA mediated enhanced leukocyte responses to infection induced after severe burn injury, it is postulated that induction of trained immunity by MyD88-activation has the potential to augment leukocyte function after trauma and restore immunocompetence in this model, a question that is currently being pursued by our group.

Our previous work demonstrated that neutrophils are also essential for MPLA-mediated infection resistance (^13^). Thus, it is possible that TLR-trained macrophages secrete chemokines for neutrophil expansion and/or mobilization and that this relationship is key for bacterial clearance and infection resistance, which remains an interesting line of future investigation. Another point to explore regarding TLR-mediated infection resistance is the potential role of organ protection and augmented efferocytosis, as the degree of bacterial clearance did not correlate with the magnitude of survival benefit and also Poly I:C-treated mice showed a trend, towards reduced bacterial burden. Regarding the later point, we postulate that slight cross-activation of downstream MyD88-signaling (evidenced by Poly I:C-induced p-IKK; Figure 1G) may drive bacterial clearance in Poly I:C-treated mice.

Our lab and others, have found that augmentation of immunometabolism is a key feature of trained immunity (^5, 23, 24^). Aerobic glycolysis, in which glucose is metabolized to lactate despite oxygen availability (*i.e.* Warburg effect) is a hallmark of trained monocytes (^25^). Ramping up glycolysis increases energy availability to support leukocyte expansion and antimicrobial function as well as generation of essential precursors for synthesis of amino acids and nucleotides necessary for proliferation and cell viability under stress conditions (^26^). Further, cellular metabolism is tightly linked with epigenetic reprogramming that occurs in trained immunity (^24^). Our previous work showed that BMDMs trained with MPLA or CpG have sustained increases in both glycolysis and mitochondrial respiration (^12, 13^). The current study builds on this work by assessing acute metabolic responses to these agonists (TLR9/MyD88-activating CpG and TLR3/TRIF-activating Poly I:C alongside dual-pathway activating MPLA) in relationship to the persistent alterations. We also employed a MyD88 or TRIF pathway knockout approach to further determine the roles of these signaling cascades. Neither glycolysis nor mitochondrial respiration were augmented by MPLA or CpG in MyD88-KO BMDMs; however, TRIF-deficient BMDMs were metabolically rewired similarly to wild type cells. Transcriptomic analysis supported these functional analyses as metabolism-related genes were similarly regulated by MPLA in WT and TRIF-deficient, but not MyD88-deficient, BMDMs.

Both transcriptionally and functionally, we found that MPLA and CpG, but not Poly I:C, augmented phagocytosis and ROS production in WT and TRIF-KO BMDMs, but not MyD88-KO BMDMs. We have previously demonstrated a direct link between metabolism and antimicrobial functions by showing that inhibition of MPLA-induced glycolysis using 2-deoxyglucose prevented enhancement of phagocytosis and chemokine secretion (^12, 13^).

Insight into underlying mechanisms of trained immunity have mostly been reported for fungal-derived β-glucan particles which also have been found to mediate host protection against a broad array of pathogens (^27–29^). Interestingly, β-glucan-mediated immunomodulation of BMDMs is dependent on the cooperation between Dectin-1 and the MyD88-activating receptor TLR2 (^30^), further implicating the role of MyD88 in trained immunity. Beta-glucan-mediated trained immunity in monocytes is characterized by aerobic glycolysis driven by a pathway involving AKT, mammalian target of rapamycin (mTOR), and hypoxia-inducible factor 1α (HIF-1α) (^24, 31^). Our previous studies have shown that TLR4-induced training of BMDMs likewise depends on mTOR and HIF-1α activation (^13^). Here, we show that the MyD88-stimulating agonist CpG activates mTOR similarly to MPLA, and further that TLR/MyD88-activation of mTOR is at least partially responsible for metabolic reprogramming and augmentation of macrophage antimicrobial function (Figure 7).

This report also builds on previous work by demonstrating that macrophage metabolic and functional programs induced by TLR/MyD88 stimulation occur in vivo. To the best of our knowledge, to date such investigations were limited to in vitro analyses, which has provided mechanistic insight but lacks physiological context. By harvesting splenic macrophages from TLR agonist-treated mice, we found that markers of TLR-induced metabolic reprogramming (mitochondrial content and intracellular ATP content) showed a similar pattern of elevation at 1dp- and 3dp-in vivo treatment (Figure 8) compared to 24h and 3dp BMDMs (Figure 4). These markers remained significantly elevated one week following CpG administration. CpG-induced enhancement of antimicrobial functional capacity (phagocytosis and ROS production) occurred in parallel. These results demonstrate that tissue-resident macrophages are reprogrammed similarly to monocyte-derived BMDMs. Interestingly, these trends correlate with the timeline of infection resistance which also began to wane at 2 weeks post-treatment (Figure 1).

Finally, we substantiated our findings using human monocyte-derived macrophages (hMDMs). Using this model, we verified that TLR/MyD88-induction of trained immunity, characterized in our model by augmentation of metabolism and antimicrobial functions, was induced by MPLA and CpG, but not Poly I:C (Figure 9). Importantly, these data indicate that our current mechanistic findings are clinically translational, and further corroborate the potential application of TLR agonists as inducers of trained immunity and therefore hold high potential as immunotherapeutic agents to augment host resistance against infections.

It is important to note that TRIF signaling may be important in trained immunity-mediated protection against viral pathogens. Previous studies have found that stimulation of TLR3 with Poly I:C provides protection against viral infections such as HSV-1(^32^) and influenza(^33^). This is plausible given that activation of TRIF signaling results in production of type I interferons which inhibit viral growth. On the other hand, other research groups have demonstrated host resistance to bacterial and viral infections via stimulation of MyD88 through TLR2 agonists MALP-2 (protection against *S. pneumoniae* (^34^)) or Pam2Cys (Influenza A (^35^)) or TLR9 agonist CpG (*L. major* (^36^), HSV-2 (^37^), MRSA(^38^)), demonstrating that activation of MyD88-signaling may confer broad protection which may extend to viral pathogens.

Together, these data show that the MyD88-signaling is critical in TLR-mediated trained immunity, which is driven, in part, by metabolic and functional reprogramming of both murine-derived and human-derived macrophages. Identification of MyD88-signaling as a key driver of trained immunity unveils more TLR ligands that may be explored for therapeutic translation. Further identification of key signaling events will continue to strengthen the prospect for therapeutic translation by potentially diminishing activation of inflammation and restoring leukocyte functions in immunosuppressed individuals. With the continued threat of antimicrobial resistance, the mission to protect vulnerable populations continues to become more urgent every day. TLR/MyD88-activating agonists such as MPLA and CpG are promising candidates to harness the potential of trained immunity whereby innate leukocytes are primed to mount a robust response to a broad array of pathogens, allowing for containment of a local infection and elimination prior to development of life-threatening sepsis.

## METHODS

### Animal model of infection and assessment of bacterial burden

#### Mice

Male BALB/c mice aged 9- to 10-weeks old were purchased from Charles River Laboratories for the majority of in vivo infection studies. MyD88^-/-^, or TRIF^-/-^ (*i.e*. Ticam1^-/-^) and the respective wild type C57BL/6J control mice were purchased from The Jackson Laboratory for select infection studies and all bone marrow-derived macrophage (BMDM) experiments. Mice were acclimated for at least one week prior to beginning experimental procedures.

#### Toll-like Receptor (TLR) agonist administration

Monophosphoryl Lipid A (MPLA) derived from *Salmonella enterica* serotype Minnesota R595 was purchased from InvivoGen and was solubilized in water containing 0.2% triethylamine and sonicated for 1 hour at 40 °C. Synthetic CpG (CpG-ODN 1826 used for murine studies, CpG-ODN 2336 used for human monocyte-derived macrophage studies), Poly I:C, and Pam3CSK4 were purchased from InvivoGen and solubilized in sterile endotoxin-free water. For in vivo treatment, TLR agonists were diluted in sterile Lactated Ringer’s (LR) solution and 20 μg (*i.v.*) was adminitered. Animals were treated for two consecutive days and infected (*S. aureus, i.v.*) the day after the 2^nd^ TLR agonist treatment. In a select experiment to the longevity of CpG-mediated resistance to infection, cohorts of animals were also infected 1- and 2-weeks after the 2^nd^ dose of agonist treatment.

#### S. aureus systemic infection

*S. aureus* obtained from American Type Culture Collection (ATCC 25923) was grown in tryptic soy broth overnight (∼22 hours) at 37 °C, centrifuged, and resuspended in sterile saline. Mice were inoculated with *S. aureus* (8 × 10^7^ – 3× 10^8^ CFU (*i.v.*). Survival, body condition, and rectal temperature were monitored at least twice daily for the first week post-infection, and at least once daily thereafter. Bacteremia was assessed by culture of small blood samples (≤10 μL, obtained by micropuncture of the tail vein using aseptic technique) diluted in sterile saline. To assess bacterial burden in the tissues, a cohort of animals were euthanized 3 days after infection, homogenates of the lung and kidney were prepared, serially diluted, and cultured on tryptic soy agar overnight (37 °C). For survival studies, animals were monitored at least daily for 14-15 days.

### Macrophage depletion

For macrophage depletion experiments, animals received 0.2 mL Clophosome-A anionic liposomes (*i.e.* clodronate liposomes) or PBS anionic control liposomes (*i.v.*, FormuMax) 1 day prior to the first dose of TLR agonist treatment.

### Immunohistochemistry

Twenty-four hours after inoculation with *S. aureus*, kidneys were harvested, fixed in 10% neutral-buffered formalin, paraffin embedded, and sectioned (5 μm). After deparaffinization, enzymatic-induced antigen retrieval was performed using Proteinase K (#S3020, Agilent Dako) for 15 minutes. The slides were incubated for 1 hour with anti-F4/80 primary antibody (1:900 dilution; NB600-404 Novus Biologicals LLC), followed by 15minute secondary antibody incubation (1:7500 dilution; rabbit anti-rat, BA-4001 Vector Laboratories, Inc). Samples from three biological replicates per experimental group were processed, and three random non-overlapping images were captured using an Olympus BX43 microscope outfitted with an Olympus DP73 camera and Olympus cellSens Dimension 1.18 software at low magnification (10X) for quantification purposes (**Supplemental Figure 1A**). A blinded investigator quantified percent area positive for F4/80 using ImageJ version 1.53e. Representative threshold-processed images are provided (**Supplemental Figure 1B**). The mean percent area of three 10X images was calculated for each biological replicate. For enhanced visualization, representative images at higher magnification (40X) were also captured.

### Bone marrow-derived macrophage (BMDM) model

Bone marrow cells were harvested and differentiated as previously described (^12, 13^). Briefly, femurs were dissected from mice and flushed with cold media (RPMI 1640 supplemented with glutamine, 25 mM HEPES, 10% fetal bovine serum, 1% antibiotic-antimycotic, and 10 ng/mL mouse recombinant M-CSF). Cells were seeded at 4 × 10^4^ cells/mL which were differentiated for 7 days as has previously been shown to yield mature macrophages (^39^). After differentiation, media was changed and cells were treated with respective TLR agonists (1 μg/mL for MPLA, CpG, and Pam3CSK, or 10 μg/mL for Poly I:C) for 24 hours or left unstimulated as a negative control. Treatment concentrations were determined based on previous studies which identified doses that stimulate appropriate induction of an inflammatory response in BMDMs (^40, 41^). After 24 hours, cells were washed with PBS and assessed immediately or rested for 3 days before assessment. When necessary, cells were replated and rested at least 1 hour prior to TLR agonist treatment. For a set of experiments, rapamycin (#13346, Cayman Chemical) was reconstituted in DMSO. Cells were treated with 100 nM rapamycin or vehicle (DMSO) 1 hour prior treatment with TLR agonists. Rapamycin remained in culture media throughout the incubation with TLR agonists until wash-off at 24h.

### Macrophage adoptive transfer

Differentiated BMDMs were treated for 24 hours with 1 μg/mL MPLA, 1 μg/mL CpG, 10 μg/mL Poly I:C, or left unstimulated. Cells were washed with PBS, lifted, resuspended in PBS, and administered to recipient mice (5.0 × 10^5^ cells/mouse) alongside vehicle control mice. Twenty-four hours later, mice were challenged with *S. aureus* (1-3X 10^8^ CFU, *i.v.*).

### Western blot

BMDMs were washed and lysed with RIPA buffer containing cOmplete Protease Inhibitor and PhosSTOP (Roche Diagnostics). Protein concentration was quantified using the BCA assay according to the manufacturer’s protocol (Bio-Rad). Samples were resolved on 4-20% Tris-glycine precast gels, and proteins were transferred to nitrocellulose membranes overnight at 4°C. Membranes were blocked with 5% fraction V BSA for 1 hour, incubated with primary antibodies overnight at 4 °C, and with secondary antibodies for 1 hour at room temperature. Protein bands were detected by chemiluminescence (ECL) and film exposure. Densitometry analysis was performed using ImageJ software (version 1.53e).

### RNA extraction and assessment of gene expression

#### RNA extraction

BMDMs were lysed using QIAshredder homogenizers and RNA was isolated using the RNeasy Mini Kit and treated with DNase (Qiagen). RNA concentration and quality were assessed preliminarily using a NanoDrop 2000 spectrophotometer (Thermo Scientific). For RNA sequencing, quality was assessed with Qubit and 2100 Bioanalyzer (Agilent Technologies) and RNA integrity number (RIN) was determined for each sample. All samples had a RIN score ≥ 7.

#### RNA sequencing and analysis

Sequencing was performed by the Vanderbilt Technologies for Advanced Genomics (VANTAGE) core facility. mRNA libraries were prepared using NEBNext Poly(A) selection (New England Biolabs). Sequencing was performed at paired-end 150 bp on an Illumina NovaSeq 6000 with at least 50 million reads per sample (average 74 million reads). RNASeq raw data in FASTQ format were analyzed by Novogene Co. Low quality reads as well as reads containing adapter and poly-N (N>10%) sequences were cleaned. Cleaned paired-end reads were aligned to the reference genome (*Mus Musculus* mm10 genome) using Spliced Transcripts Alignment to a Reference (STAR) software. FeatureCounts was used to count the read numbers mapped of each gene and reads per kilobase of exon model per million mapped reads (RPKM) were calculated. Differential expression analysis between two groups was performed using DESeq2 R package which uses a model based on the negative binomial distribution. The resulting *p* values were adjusted using the Benjamini and Hochberg’s approach for controlling false discovery rate (FDR), and genes with an adjusted *p* < 0.05 and log2 fold change >1 were considered differentially expressed. Hierarchical cluster analysis was performed using Integrated Differential Expression and Pathway analysis (idep, version iDEP.951) on readcount data using complete linkage and correlation-based distance. Gene Ontology (GO) enrichment analysis was conducted using the clusterProfiler R package.

#### Quantitative real-time PCR

After quantification and passing quality control, RNA was reverse transcribed into cDNA using iScript cDNA Synthesis Kit (#170-8891, Bio-Rad) according to the manufacturer’s protocol. SsoFast EvaGGreen Supermix (#1725200, Bio-Rad) was used to perform quantitative real-time PCR (qRT-PCR) which was performed on a Bio-Rad CFX96 housed in the Molecular Cell Biology Resource Core at Vanderbilt University Medical Center. Gene expression was normalized to beta-actin as an endogenous control, and fold change was calculated by the comparative ΔΔCT method. Primer sequences for the target genes are provided in **Supplemental Table 1**.

### Assessment of macrophage metabolism

#### Seahorse extracellular flux analysis

BMDMs were plated at 5X10^4^ cells/well with at least 3 technical replicates per group. Glycolysis and mitochondrial stress tests were performed according to the manufacturer’s protocol using a Seahorse XF^e^96 Extracellular Flux Analyzer (Agilent Technologies). Glycolytic capacity was measured by extracellular acidification rate (ECAR) at baseline and after sequential treatment with 10 mM glucose, 1 μM oligomycin, and 50 mM 2-deoxyglucose. Mitochondrial respiration was measured by oxygen consumption rate (OCR) at baseline and after sequential treatment with 1 μM oligomycin, 1 μM FCCP, and 0.5 μM antimycin A and rotenone. The measurements were corrected for background, and technical replicates were averaged. Of note, the Seahorse XF^e^96 Extracellular Flux Analyzer was serviced between sets of experiments and the subsequent ECAR data obtained from the glycolysis stress test resulted in consistent trends among groups but reduced overall scale.

#### ATP quantification

Macrophages (5X10^5^ cells) were lysed with Luciferase Cell Culture Lysis Reagent (Promega). A commercially available ATP Determination Kit (Invitrogen) was used to and the resulting luminescence of at least two technical replicates per sample was measured using a BioTek Synergy MX plate reader.

#### MitoTracker and TMRM staining

For BMDMs, 2.5X10^5^ cells were plated in a 24-well plate one day prior to the assays. Cells were washed with phenol red-free RPMI and incubated with MitoTracker green (50 nM, Invitrogen) or TMRM dye (100 nM, Invitrogen) alongside unstained controls for 30 minutes at 37 °C. For isolated splenic macrophages, at least 5X10^4^ cells were suspended in the respective dyes and incubated for 30 minutes at 37 °C. Following the incubation, cells were washed, and MFI was measured using an Accuri C6 flow cytometer. Data was analyzed using Accuri C6 software (BD Biosciences).

### Assays of antimicrobial functions

#### Phagocytosis

BMDMs were plated in a black well/clear bottom 96-well plate (5 × 10^4^ cells/well) at least in triplicate per group. pHrodo-tagged red *S. aureus* bioparticles were resuspended in phenol red-free RPMI and added (100 μg) to washed cells, incubated for 1 hour, and fluorescence was measured every 15-minutes on a BioTek Synergy MX plate reader.

#### Bacterial killing

BMDMs were plated at 1X10^5^ cells/well of a 96-well plate (3-5 replicates per group). Cells were washed with antibiotic-free media with 0.1% FBS 5 times and replaced with 100 μL of media containing *S. aureus* at a multiplicity of infection (MOI) of 10. Cells were incubated with the inoculum for 1 hour at 37 °C. The media was collected for determination of extracellular bacteria. Macrophages were washed with PBS and remaining extracellular bacteria were killed with media containing 300 μg/mL gentamicin and 0.1%FBS. The BMDMs were washed in triplicate and lysed. Samples were serially diluted and cultured overnight for determination of CFU.

#### ROS production

ROS production by BMDMs (1X10^5^ events) was assessed using a commercially available kit (Respiratory Burst Assay, Cayman Chemical) according to the manufacturer’s protocol with the exception of stimulating the cells with PMA. Thus, incubating with dihydrorodamine 123 was an indicator of baseline ROS production, which was quantified by flow cytometry.

### Splenic macrophage isolation

Mice were treated with CpG (20 μg *i.v.*) for two consecutive days alongside vehicle treated controls, and spleens were harvested 1 day, 3 days, 1 week, or 2 weeks after the 2^nd^ treatment. Spleens were transferred to a pre-wet cell strainer (70 μm) set in a 50-mL tube pre-coated with Sigmacote (Sigma Aldrich). Spleens were crushed using the flat end of a syringe plunger, and 20 mL of 50% Accumax (Sigma Aldrich) in dPBS was passed through the filter. Cells were centrifuged at 300 X g for 10 minutes at 4 °C. The supernatant was decanted, and the pellet was resuspended in EasySep cell separation buffer (2% FBS, 1 mM EDTA in dPBS). The cell suspension was passed through a cell strainer before counting. Samples were pelleted and resuspended in separation buffer at 1 × 10^8^ cells/mL.

Splenic macrophages were magnetically labeled and isolated using StemCell EasySep PE Positive Selection Kit II in concert with an F4/80 PE-conjugated antibody (StemCell Technologies). Briefly, the cell suspensions were incubated with F4/80-PE (3 μg/mL) for 15 minutes at room temperature. Cells were washed, incubated with the selection cocktail (15 minutes), and then with the dextran-coated magnetic particles (RapidSpheres, 10 minutes). Tubes were placed into an EasyEights magnet for 10 minutes, after which the supernatant was removed with a serological pipette. The cells were resuspended in isolation buffer and incubated for 5 minutes in the magnet, which was repeated three additional times, yielding >95% viable, >85% pure F4/80^+^ macrophages (**Supplemental Figure 8**) which were used immediately for downstream assays. Positive magnetic isolation allows for increased yield and purity of cell populations (^42^). Alternatively, plastic adhesion has been shown to result in low yield and viability (^43^), while overlap of markers prevented the use of negative isolation. Importantly, no differences in gene expression of IL-6, TNF-α, or CD206 have been reported to be observed in cell populations isolated by these three methods (^43^).

### Human monocyte-derived macrophage (hMDM) model

#### Isolation of monocytes from human buffy coat samples

Fresh buffy coat samples collected in EDTA from healthy male donors (age range 22-56; 14% Caucasian, 57% Hispanic, and 14% Black) were purchased from BioIVT. Samples were collected under IRB-approved protocols and were shipped the same day of collection. Mononuclear cells (MNCs) were enriched using Lymphoprep density gradient medium (Catalog #07811) and SepMate tubes (Catalog #85450; STEMCELL Technologies). Briefly, buffy coat samples were diluted 1:1 in PBS containing 2% FBS and the recommended volume was added to SepMate tubes containing 15mL room temperature Lymphoprep medium. Tubes were centrifuged at 1,200X*g* for 20 minutes at room temperature, the plasma layer was pipetted off, and the MNC layer was poured into a fresh tube.

Cells were washed, counted, and resuspended in the recommended buffer (1mM EDTA and 2% FBS in PBS) at 1X10^8^ cells/mL. CD14^+^ monocytes were magnetically isolated (#100-0694 EasySep human positive selection kit II, STEMCELL Technologies) and the resulting cell suspension was passed through a 70 μm cell strainer.

#### Macrophage differentiation and training

Cells were seeded at 4X10^4^ cells/mL in RPMI 1640 media supplemented with glutamine, 25 mM HEPES, 10% fetal bovine serum, 100 units/100 μg penicillin-streptomycin, and 25 ng/mL human recombinant M-CSF (#216-MC, R&D Systems Inc). Cells were differentiated for 7-10 days prior to treatment as has previously been shown to yield mature macrophages (^44^). Human monocyte-derived macrophages (hMDMs) were treated with 10 μg/mL MPLA, 10 μg/mL CpG, or 100 μg/mL Poly I:C based on our previous work (^45^). In alignment with our BMDM studies, hMDMs were treated for 24h with the respective agonists before wash-off, and then rested for 3 days prior to assays.

#### hMDM assays

Seahorse extracellular flux analysis was performed as described above for BMDMs with the following modifications: hMDMs were replated 6h prior to conducting the assays, and plated at 8X10^4^ cells/well and 2X10^4^ cells/well for glycolysis stress test and mitochondrial stress test, respectively. Mitotracker and TMRM staining were conducted as described except cells were used at 4X10^4^ cells/well of a 6-well plate. Phagocytosis was assayed by suspending 5X10^5^ hMDMs with 100 μL of pHrodo-tagged red S. aureus bioparticles which was incubated for 2 hours and MFI was measured by flow cytometry. ROS production was performed as described above.

### Statistical analysis

All data were analyzed using GraphPad Prism 8. Survival data were analyzed using Kaplan-Meier curves with a Log Rank test to detect statistical significances between groups. Student’s t-test was performed for comparison of two groups. One-way ANOVA analysis was performed for comparisons of multiple groups followed by Dunnett’s post hoc multiple comparison test. In experiments where data were not normally distributed, we employed non-parametric statistics. A *p* value of ≤ 0.05 was considered statistically significant. Differences in gene expression were determined as described above. Data are expressed as mean ± SEM unless otherwise noted.

### Study approval

All animal experimental procedures were performed in accordance with the National Institutes of Health (NIH) guidelines for ethical treatment and were approved by the Institutional Animal Care and Use Committee (IACUC) at Vanderbilt University Medical Center (Protocol ID# M-1800068-01).

## AUTHOR CONTRIBUTIONS

All authors provided critical feedback and helped design the research. AMO, LL, CLS, JFS, ERS, and JKB conceived and planned the experiments. AMO, LL, CLS, KRB, MAM, JFS, TKP, and JKB carried out experiments. AMO, CLS, KRB, MAM, and TKP contributed to sample preparation. JW provided project management and coordination. OB and KS helped curate data. AH, NKP, ERS, and JKB provided experimental expertise, supervision, and assisted with data interpretation. AMO, ERS, and JKB wrote the manuscript. All authors helped edit the manuscript and have approved the final version.

## DATA AVAILABILITY

The data that support the findings of this study are available in Gene Expression Omnibus (GEO) under accession number GSE205186.

## ACKNOWLEDGEMENTS

This work was supported by the National Institutes of Health (NIH) Grants R01 GM121711 and R35 GM141927 awarded to JKB, R01 AI151210 and R01 GM119197 awarded to ERS, T32AI38932-02 and VICTR Voucher #VR56306 awarded to AMO, K08 GM123345 awarded to AH, T32 GM108554 awarded to NKP, who was also supported by a Vanderbilt Faculty Research Scholars Award and Shock Society Faculty Research Award. This study utilized the Agilent Seahorse Extracellular Flux Analyzer that is maintained within the Vanderbilt High-Throughput Screening Core Facility (supported by NIH Shared Instrumentation Grant 1S10OD018015). We thank the Translational Pathology Shared Resource for performing tissue processing and immunohistochemical analysis (supported by NCI/NIH Cancer Center Support Grant 5P30 CA68485-1 and The Shared Instrumentation Grant S10 OD023475-01A1), and the Vanderbilt Technologies for Advanced Genomics (VANTAGE) core facility for performing RNASeq. Schematic illustrations were created with BioRender.com and publication licenses were obtained.

## Notes

### Competing Interest Statement

The authors have declared no competing interest.

