## Supplemental Table 1 and Supplemental Figures 1-8 for "MyD88-Dependent Signaling Drives Toll-Like Receptor-Induced Trained Immunity in Macrophages"

**<u>Table 1</u>. Primer sequences used for qRT-PCR gene expression analyses.**

| Gene | Forward primer (5’→3’) | Reverse primer (5’→3’) |
| --- | --- | --- |
| *MARCO* | CCTCCAGGGACTTACGGGT | CCAGTGAGACCTATGTCACCT |
| *PECAM1* | ACGCTGGTGCTCTATGCAAG | TCAGTTGCTGCCCATTCATCA |
| *Ptprm* | CATGCTGGTCAACACTTCTGG | GCAGCGTTACTCTTGCTGGATA |

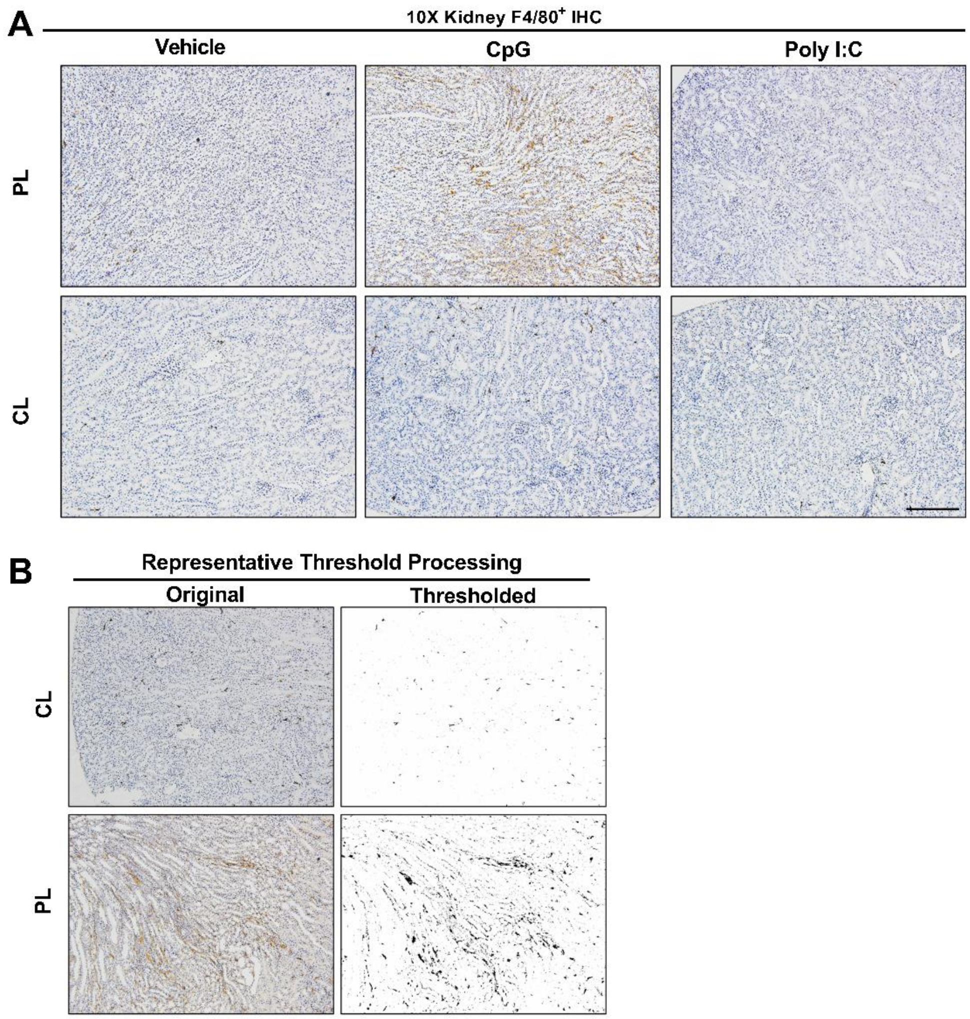


**<u>Supplemental Figure 1</u>. F4/80 immunohistochemical staining of kidney samples harvested 1-day after *S. aureus* infection in macrophage-depleted mice compared to controls.** (**A**) Representative low magnification (10X) micrographs were captured (scale bar represents 100 μm) and (**B**) threshold processing was conducted for quantification using ImageJ in clodronate liposome (CL)-treated or PBS liposome (PL)-treated mice administered TLR agonists CpG or Poly I:C (20 μg) alongside vehicle controls (0.2 mL *i.v.*).

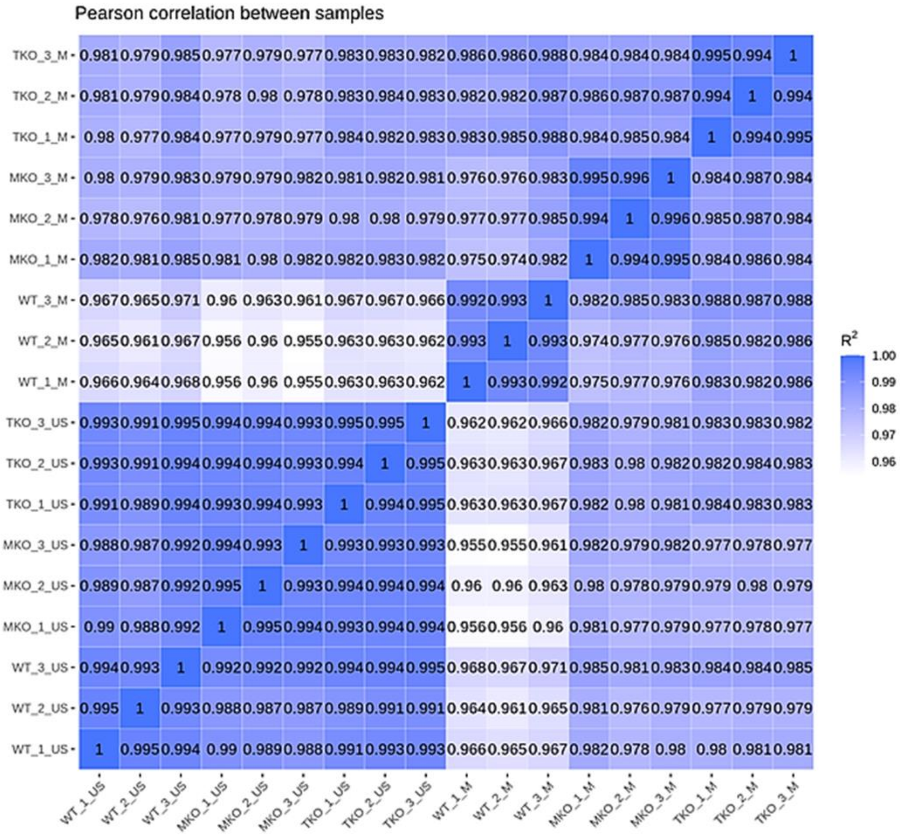


**<u>Supplemental Figure 2</u>. Pearson correlation analysis demonstrates high consistency among biological replicates.** Pearson correlation analysis on WT, MyD88 KO (MKO), or TRIF KO (TKO) RNA samples from unstimulated (US) or 3dp MPLA (M) bone marrow derived macrophage conditions demonstrates the degree of correlation ($R^2$) among biological replicates (n = 3/group). Darker shading and higher $R^2$ value indicate stronger linear correlation.

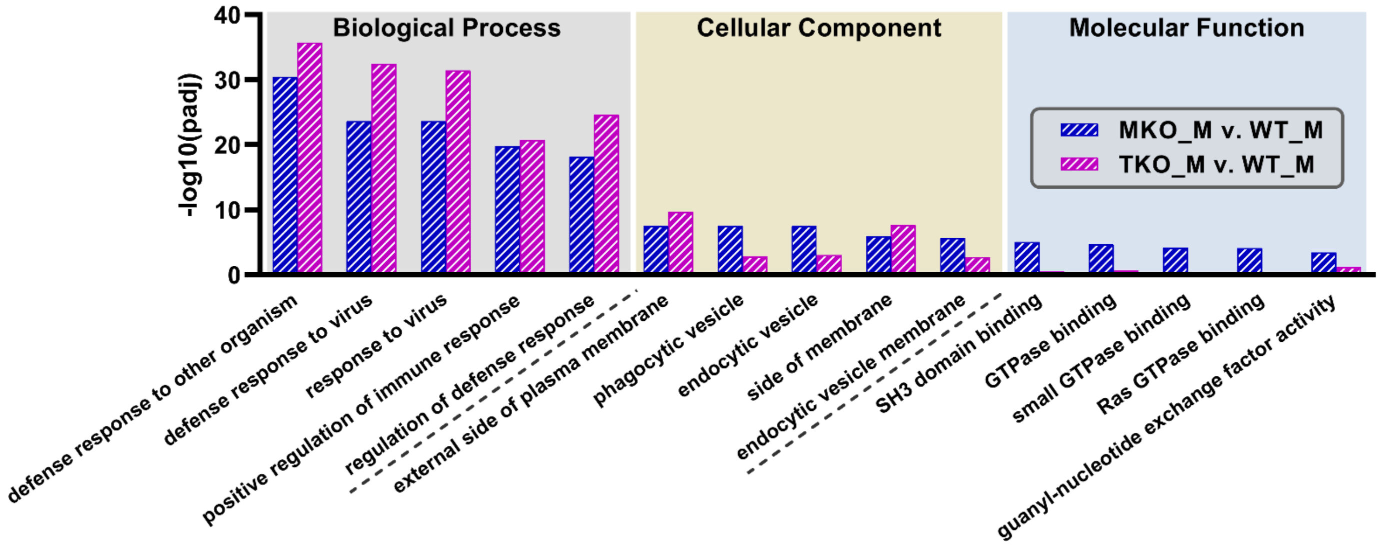


**<u>Supplemental Figure 3</u>. Downregulated enrichment pathways in MPLA-treated MyD88 KO (MKO) or TRIF KO (TKO) compared to MPLA-treated wild type (WT) macrophages.** The 5 most downregulated enrichment pathways of each gene ontology (GO) class (biological process, cellular component, and molecular function) in MPLA-treated MKO cells compared to MPLA-treated WT cells were identified. These identified pathways were then assessed for MPLA-treated TKO cells compared to MPLA-treated WT macrophages, and the degree of significance is displayed (-log padj).

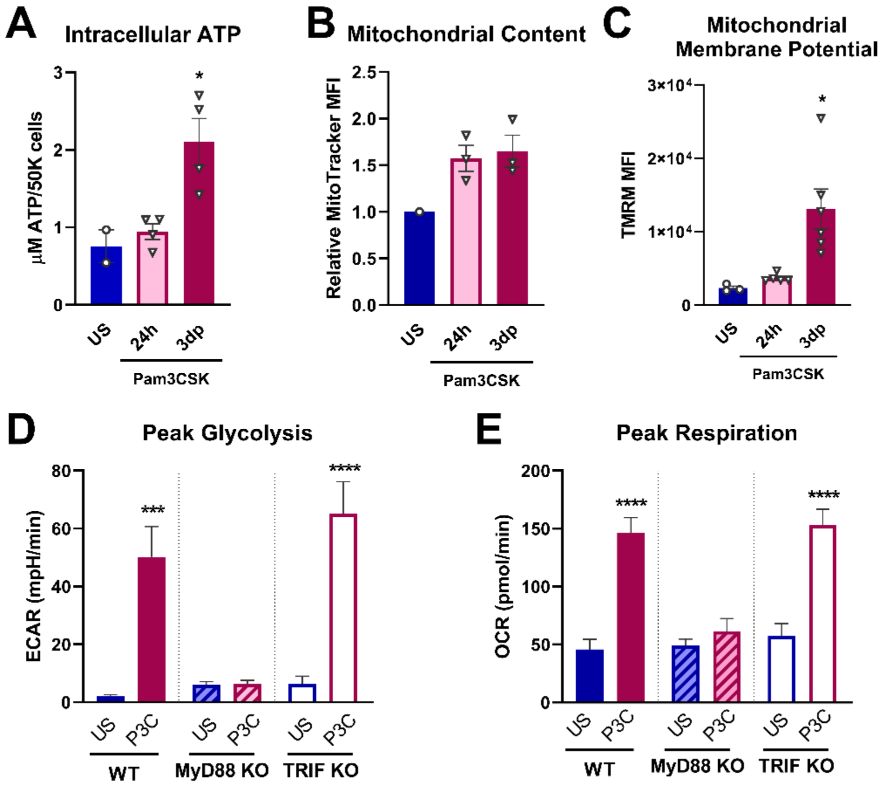


**<u>Supplemental Figure 4</u>. TLR2/MyD88-activating agonist Pam3CSK triggers macrophage metabolic reprogramming.** Bone marrow-derived macrophages (BMDMs) were treated with Pam3CSK (P3C; 1 µg/mL) for 24 hours and allowed to rest for 3 days compared to unstimulated (US) controls. (**A**) Intracellular ATP was measured. (**B**) Mitochondrial content was assessed by MitoTracker Green staining and (**C**) mitochondrial membrane potential was assessed by TMRM staining and measured by flow cytometry with mean fluorescent intensity (MFI) shown. (**D**) BMDMs from wild type (WT), MyD88 KO, and TRIF KO mice were treated with P3C as described and compared to US controls. Peak glycolysis was measured as indicated by Extracellular Acidification Rate (ECAR), and (**E**) peak mitochondrial respiration was measured by oxygen consumption rate (OCR). Data are shown as mean ± SEM, n = 3-5/group, * $p < 0.05$, *** $p < 0.001$, **** $p < 0.0001$ by ANOVA followed by Dunnett's post-hoc multiple comparison test.

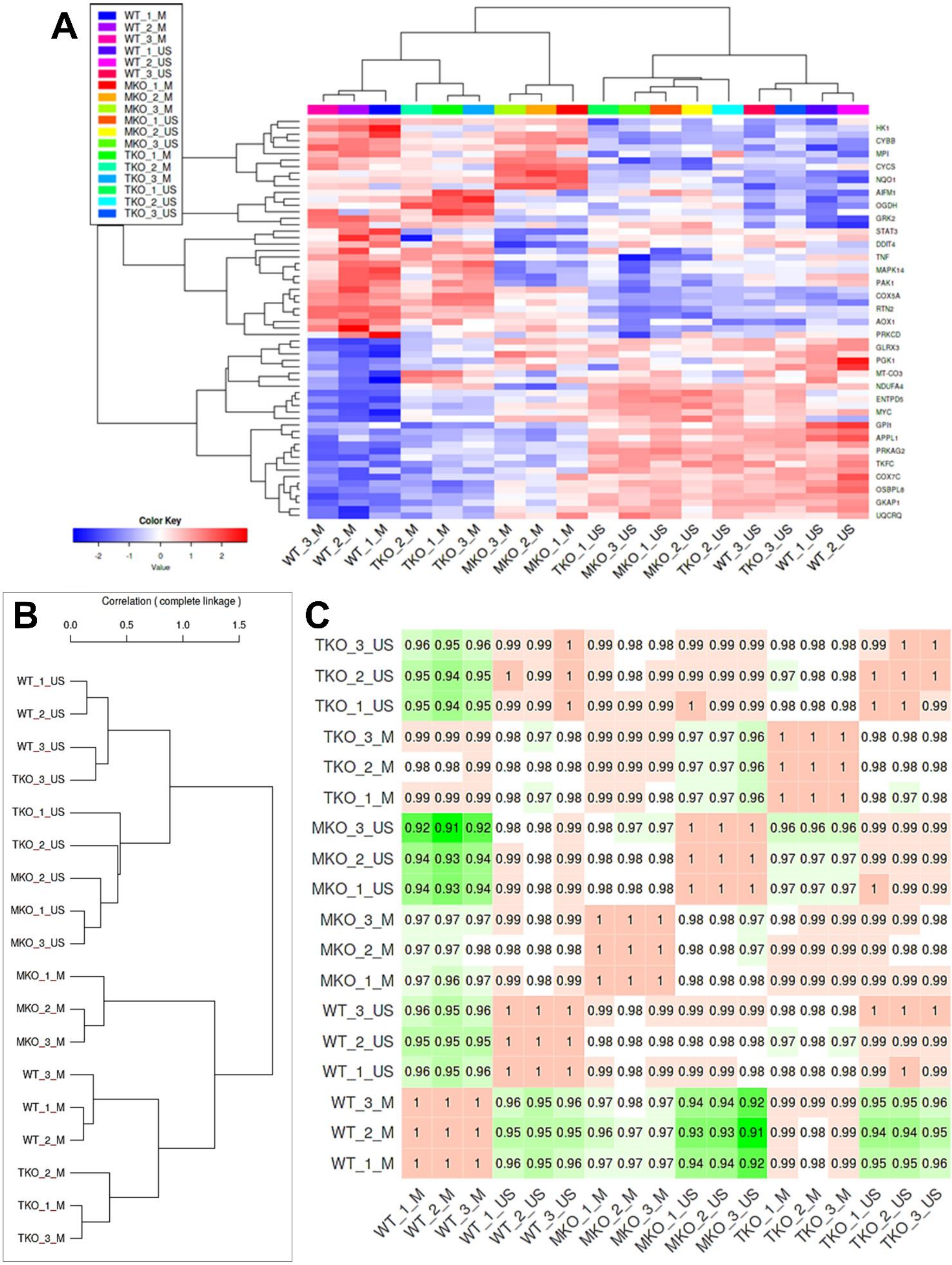
A
WT_1_M
WT_2_M
WT_3_M
WT_1_US
WT_2_US
WT_3_US
MKO_1_M
MKO_2_M
MKO_3_M
MKO_1_US
MKO_2_US
MKO_3_US
TKO_1_M
TKO_2_M
TKO_3_M
TKO_1_US
TKO_2_US
TKO_3_US
HK1
CYBB
MPI
CYCS
NQO1
AIFM1
OGDH
GRK2
STAT3
DDIT4
TNF
MAPK14
PAK1
COX5A
RTN2
AOX1
PRKCD
GLRX3
PGK1
MT-CO3
NDUFA4
ENTPD5
MYC
GPI1
APPL1
PRKAG2
TKFC
COX7C
OSBPL8
GKAP1
UQCRQ
Color Key
-2 -1 0 1 2
Value
WT_3_M WT_2_M WT_1_M TKO_2_M TKO_1_M TKO_3_M MKO_3_M MKO_2_M MKO_1_M TKO_1_US MKO_3_US MKO_1_US MKO_2_US TKO_2_US WT_3_US TKO_3_US WT_1_US WT_2_US
B
Correlation ( complete linkage )
0.0 0.5 1.0 1.5
WT_1_US
WT_2_US
WT_3_US
TKO_3_US
TKO_1_US
TKO_2_US
MKO_2_US
MKO_1_US
MKO_3_US
MKO_1_M
MKO_2_M
MKO_3_M
WT_3_M
WT_1_M
WT_2_M
TKO_2_M
TKO_1_M
TKO_3_M
C
TKO_3_US 0.96 0.95 0.96 0.99 0.99 1 0.99 0.98 0.98 0.99 0.99 0.99 0.98 0.98 0.98 0.99 1 1
TKO_2_US 0.95 0.94 0.95 1 0.99 1 0.99 0.98 0.99 0.99 0.99 0.99 0.97 0.98 0.98 1 1 1
TKO_1_US 0.95 0.94 0.95 0.99 0.99 1 0.99 0.99 0.99 1 0.99 0.99 0.98 0.98 0.98 1 1 0.99
TKO_3_M 0.99 0.99 0.99 0.98 0.97 0.98 0.99 0.99 0.99 0.97 0.97 0.96 1 1 1 0.98 0.98 0.98
TKO_2_M 0.98 0.98 0.99 0.98 0.98 0.98 0.99 0.99 0.99 0.97 0.97 0.96 1 1 1 0.98 0.98 0.98
TKO_1_M 0.99 0.99 0.99 0.98 0.97 0.98 0.99 0.99 0.98 0.97 0.97 0.96 1 1 1 0.98 0.97 0.98
MKO_3_US 0.92 0.91 0.92 0.98 0.98 0.99 0.98 0.97 0.97 1 1 1 0.96 0.96 0.96 0.99 0.99 0.99
MKO_2_US 0.94 0.93 0.94 0.99 0.98 0.99 0.98 0.98 0.98 1 1 1 0.97 0.97 0.97 0.99 0.99 0.99
MKO_1_US 0.94 0.93 0.94 0.99 0.98 0.99 0.98 0.98 0.98 1 1 1 0.97 0.97 0.97 1 0.99 0.99
MKO_3_M 0.97 0.97 0.97 0.99 0.98 0.99 1 1 1 0.98 0.98 0.97 0.98 0.99 0.99 0.99 0.99 0.98
MKO_2_M 0.97 0.97 0.98 0.98 0.98 0.98 1 1 1 0.98 0.98 0.97 0.99 0.99 0.99 0.99 0.98 0.98
MKO_1_M 0.97 0.96 0.97 0.99 0.98 0.99 1 1 1 0.98 0.98 0.98 0.99 0.99 0.99 0.99 0.99 0.99
WT_3_US 0.96 0.95 0.96 1 1 1 0.99 0.98 0.99 0.99 0.99 0.99 0.98 0.98 0.98 1 1 1
WT_2_US 0.95 0.95 0.95 1 1 1 0.98 0.98 0.98 0.98 0.98 0.98 0.97 0.98 0.97 0.99 0.99 0.99
WT_1_US 0.96 0.95 0.96 1 1 1 0.99 0.98 0.99 0.99 0.99 0.98 0.98 0.98 0.98 0.99 1 0.99
WT_3_M 1 1 1 0.96 0.95 0.96 0.97 0.98 0.97 0.94 0.94 0.92 0.99 0.99 0.99 0.95 0.95 0.96
WT_2_M 1 1 1 0.95 0.95 0.95 0.96 0.97 0.97 0.93 0.93 0.91 0.99 0.98 0.99 0.94 0.94 0.95
WT_1_M 1 1 1 0.96 0.95 0.96 0.97 0.97 0.97 0.94 0.94 0.92 0.99 0.98 0.99 0.95 0.95 0.96
WT_1_M WT_2_M WT_3_M WT_1_US WT_2_US WT_3_US MKO_1_M MKO_2_M MKO_3_M MKO_1_US MKO_2_US MKO_3_US TKO_1_M TKO_2_M TKO_3_M TKO_1_US TKO_2_US TKO_3_US

**Supplemental Figure 5. Cluster analysis of metabolism-related genes demonstrates distance of MPLA-treated MyD88 KO macrophages from MPLA-treated wild type and TRIF KO macrophages.** WT, MKO, and TKO bone marrow-derived macrophages (BMDMs) were treated with MPLA for 24h and rested for 3 days (_M) or left unstimulated as controls (_US) and RNA was harvested for RNAseq analysis. (**A**) Normalized expression of metabolism-related genes (list provided in Figure 5A) in individual samples (represented by columns, n=3/group) represented as a heatmap. Complete-linkage hierarchical clustering was performed using correlation-based distance. (**B**) Hierarchical clustering tree (using parameters described in panel A) for genes with maximum expression level at the top 75%. (**C**) Pearson's Correlation matrix demonstrating stronger positive correlation (r value closer to 1) in red and weaker correlation in green.

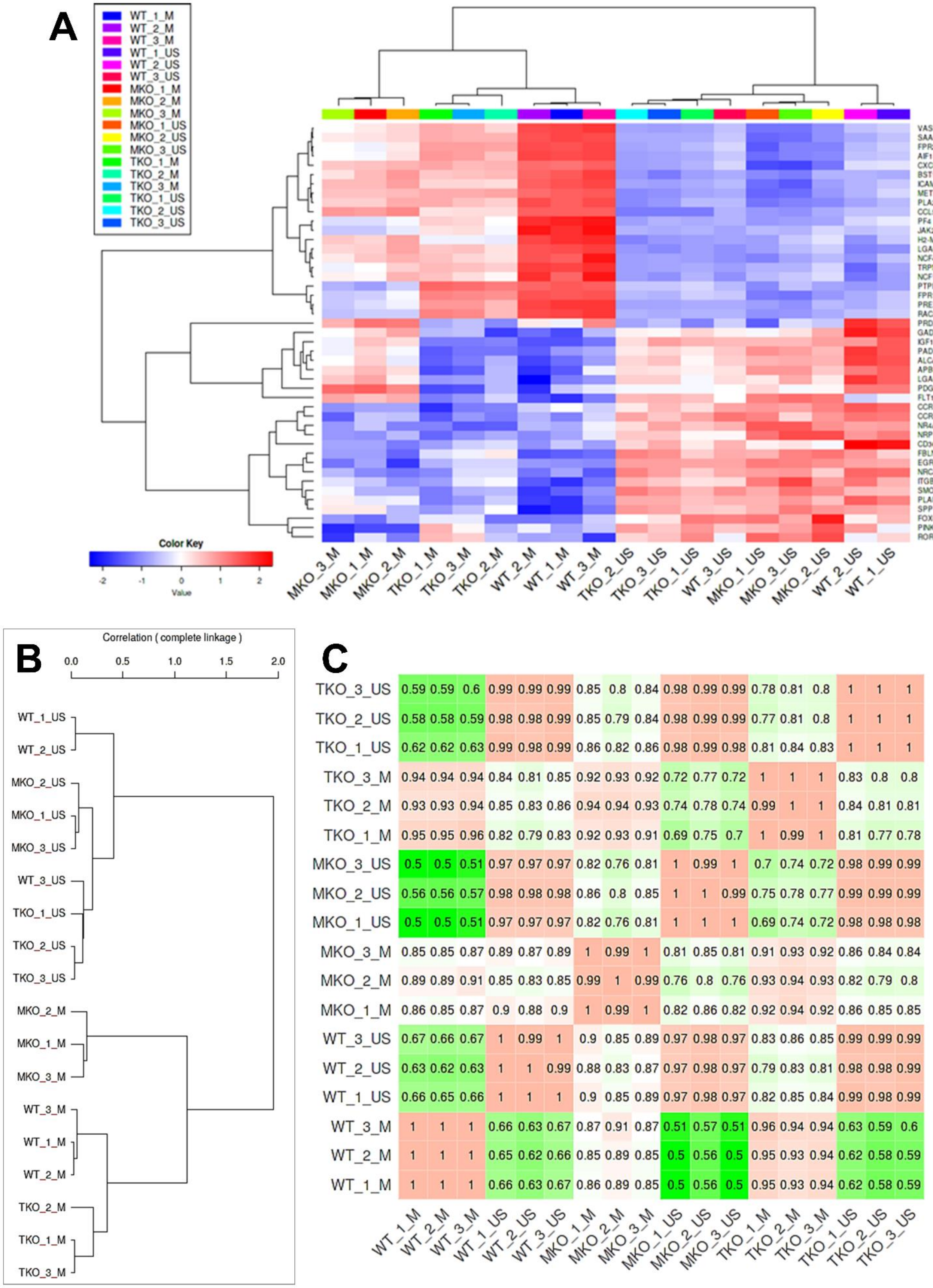
A
WT_1_M
WT_2_M
WT_3_M
WT_1_US
WT_2_US
WT_3_US
MKO_1_M
MKO_2_M
MKO_3_M
MKO_1_US
MKO_2_US
MKO_3_US
TKO_1_M
TKO_2_M
TKO_3_M
TKO_1_US
TKO_2_US
TKO_3_US
VASP
SAA3
FPR2
AIF1
CXCL16
BST1
ICAM1
MET
PLA2G7
CCL5
PF4
JAK2
H2-M3
LGALS9
NCF4
TRPM2
NCF1
PTPRM
FPR1
PREX1
RAC2
PRDX1
GADD45A
IGF1
PADI2
ALCAM
APBB2
LGALS3
PDGFB
FLT1
CCR1
CCR2
NR4A1
NRP1
CD36
FBLN5
EGR3
NRCAM
ITGB3
SMO
PLAU
SPP1
FOXO1
PINK1
RORA
Color Key
-2 -1 0 1 2
Value
MKO_3_M MKO_1_M MKO_2_M TKO_1_M TKO_3_M TKO_2_M WT_2_M WT_1_M WT_3_M TKO_2_US TKO_3_US TKO_1_US WT_3_US MKO_1_US MKO_3_US MKO_2_US WT_2_US WT_1_US
B
Correlation ( complete linkage )
0.0 0.5 1.0 1.5 2.0
WT_1_US
WT_2_US
MKO_2_US
MKO_1_US
MKO_3_US
WT_3_US
TKO_1_US
TKO_2_US
TKO_3_US
MKO_2_M
MKO_1_M
MKO_3_M
WT_3_M
WT_1_M
WT_2_M
TKO_2_M
TKO_1_M
TKO_3_M
C
TKO_3_US 0.59 0.59 0.6 0.99 0.99 0.99 0.85 0.8 0.84 0.98 0.99 0.99 0.78 0.81 0.8 1 1 1
TKO_2_US 0.58 0.58 0.59 0.98 0.98 0.99 0.85 0.79 0.84 0.98 0.99 0.99 0.77 0.81 0.8 1 1 1
TKO_1_US 0.62 0.62 0.63 0.99 0.98 0.99 0.86 0.82 0.86 0.98 0.99 0.98 0.81 0.84 0.83 1 1 1
TKO_3_M 0.94 0.94 0.94 0.84 0.81 0.85 0.92 0.93 0.92 0.72 0.77 0.72 1 1 1 0.83 0.8 0.8
TKO_2_M 0.93 0.93 0.94 0.85 0.83 0.86 0.94 0.94 0.93 0.74 0.78 0.74 0.99 1 1 0.84 0.81 0.81
TKO_1_M 0.95 0.95 0.96 0.82 0.79 0.83 0.92 0.93 0.91 0.69 0.75 0.7 1 0.99 1 0.81 0.77 0.78
MKO_3_US 0.5 0.5 0.51 0.97 0.97 0.97 0.82 0.76 0.81 1 0.99 1 0.7 0.74 0.72 0.98 0.99 0.99
MKO_2_US 0.56 0.56 0.57 0.98 0.98 0.98 0.86 0.8 0.85 1 1 0.99 0.75 0.78 0.77 0.99 0.99 0.99
MKO_1_US 0.5 0.5 0.51 0.97 0.97 0.97 0.82 0.76 0.81 1 1 1 0.69 0.74 0.72 0.98 0.98 0.98
MKO_3_M 0.85 0.85 0.87 0.89 0.87 0.89 1 0.99 1 0.81 0.85 0.81 0.91 0.93 0.92 0.86 0.84 0.84
MKO_2_M 0.89 0.89 0.91 0.85 0.83 0.85 0.99 1 0.99 0.76 0.8 0.76 0.93 0.94 0.93 0.82 0.79 0.8
MKO_1_M 0.86 0.85 0.87 0.9 0.88 0.9 1 0.99 1 0.82 0.86 0.82 0.92 0.94 0.92 0.86 0.85 0.85
WT_3_US 0.67 0.66 0.67 1 0.99 1 0.9 0.85 0.89 0.97 0.98 0.97 0.83 0.86 0.85 0.99 0.99 0.99
WT_2_US 0.63 0.62 0.63 1 1 0.99 0.88 0.83 0.87 0.97 0.98 0.97 0.79 0.83 0.81 0.98 0.98 0.99
WT_1_US 0.66 0.65 0.66 1 1 1 0.9 0.85 0.89 0.97 0.98 0.97 0.82 0.85 0.84 0.99 0.98 0.99
WT_3_M 1 1 1 0.66 0.63 0.67 0.87 0.91 0.87 0.51 0.57 0.51 0.96 0.94 0.94 0.63 0.59 0.6
WT_2_M 1 1 1 0.65 0.62 0.66 0.85 0.89 0.85 0.5 0.56 0.5 0.95 0.93 0.94 0.62 0.58 0.59
WT_1_M 1 1 1 0.66 0.63 0.67 0.86 0.89 0.85 0.5 0.56 0.5 0.95 0.93 0.94 0.62 0.58 0.59
WT_1_M WT_2_M WT_3_M WT_1_US WT_2_US WT_3_US MKO_1_M MKO_2_M MKO_3_M MKO_1_US MKO_2_US MKO_3_US TKO_1_M TKO_2_M TKO_3_M TKO_1_US TKO_2_US TKO_3_US

**Supplemental Figure 6. Cluster analysis of antimicrobial function-related genes demonstrates distance of MPLA-treated MyD88 KO macrophages from MPLA-treated wild type and TRIF KO macrophages.** WT, MKO, and TKO bone marrow-derived macrophages (BMDMs) were treated with MPLA for 24h and rested for 3 days (_M) or left unstimulated as controls (_US) and RNA was harvested for RNAseq analysis. (**A**) Normalized expression of antimicrobial function-related genes (list provided in Figure 6C) in individual samples (represented by columns, n=3/group) represented as a heatmap. Complete-linkage hierarchical clustering was performed using correlation-based distance. (**B**) Hierarchical clustering tree (using parameters described in panel A) for genes with maximum expression level at the top 75%. (**C**) Pearson's Correlation matrix demonstrating stronger positive correlation (r value closer to 1) in red and weaker correlation in green.

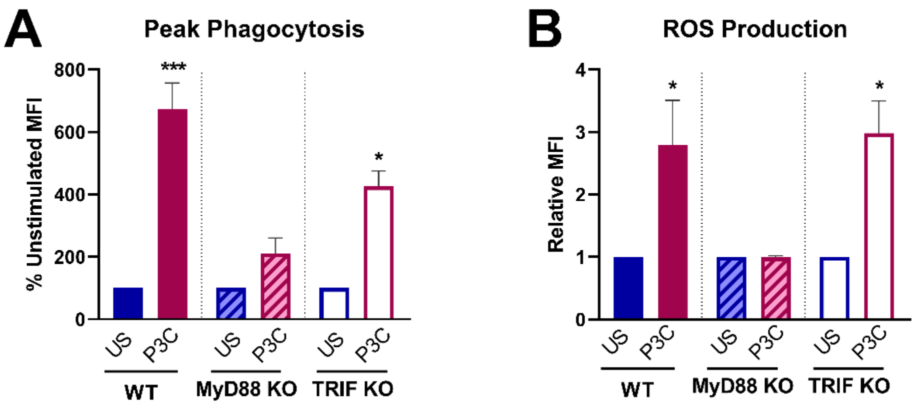


<u>Supplemental Figure 7</u>**. TLR2/MyD88-stimulating agonist Pam3CSK augments antimicrobial capacity of BMDMs.** Bone marrow-derived macrophages (BMDMs) from wild type (WT), MyD88 KO, and TRIF KO mice were treated with Pam3CSK (P3C; 1 µg/mL) for 24 hours and allowed to rest for 3 days compared to unstimulated (US) controls. (**A**) Phagocytic capacity and (**B**) reactive oxygen species (ROS) production capacity were measured.. Data are shown as mean ± SEM, n = 2-7/group, * $p < 0.05$, *** $p < 0.001$ by Student's T-test.

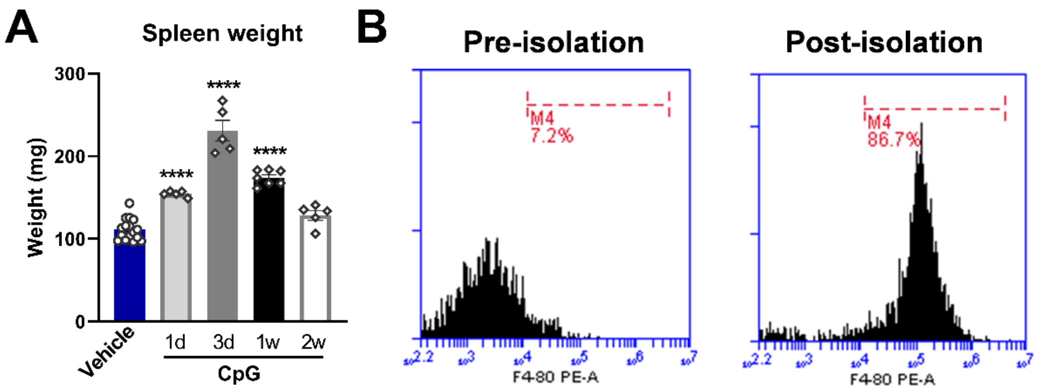


<u>Supplemental Figure 8</u>. **Splenomegaly was observed up to 1-week after CpG treatment and F4/80+ splenic macrophages were immunomagnetically isolated for downstream analysis.** (**A**) Spleen weight (mg) was measured 1d, 3d, 1-week, and 2-weeks after CpG treatment (20µg *i.v.*; n=4-16/group). (**B**) Percent F4/80$^+$ macrophages in splenocyte cell suspension before (left) and after (right) magnetic isolation as assessed by flow cytometry. **** $p < 0.0001$ by ANOVA followed by Dunnett's post-hoc multiple comparison test.
